# CYSTEINE-RICH RLK2 regulates development *via* callose synthase-dependent symplastic transport in *Arabidopsis*

**DOI:** 10.64898/2025.12.03.691776

**Authors:** Adam Zeiner, Julia Krasensky-Wrzaczek, Sunita Jindal, Yanbiao Sun, David Ušák, Jakub Hajný, Mansi Sharma, Filis Morina, Elisa Andresen, Mirva Pääkkönen, Hendrik Küpper, Johannes Merilahti, Roman Pleskot, Charles W. Melnyk, Michael Wrzaczek

## Abstract

CYSTEINE-RICH RECEPTOR-LIKE PROTEIN KINASEs (CRKs) play an important role in plant development and stress responses. One of the best described members of the Arabidopsis CRK family is CRK2, which was proposed as a crucial regulator of intercellular transport facilitated by plasmodesmata (PD). As intercellular channels allowing symplastic communication, PD-mediated transport is predominantly regulated by callose synthase (CALS)-mediated callose deposition. This process can impact not just the distribution of molecules between adjacent cells, but also the symplastic loading of vascular tissue, thereby influencing plant stress responses and developmental processes. Here we described the overlapping expression pattern of genes encoding phylogenetically closely related CALS1 and CALS3. Both CALSs were phosphorylated *in vitro* by CRK2, and the genetic interaction between genes encoding CRK2 and CALS1 or CALS3 revealed their impact on callose deposition, rosette growth, primary root length, and development, represented as a decreased number of true leaves. Importantly, we observed significant accumulation of starch in *crk2* mutant plants, especially in developmentally older leaves, which was reverted by the independent introduction of *cals1.5* and *cals3.1* into the *crk2* mutant background. The observed starch accumulation was accompanied by photosynthesis inhibition. We propose that the growth and developmental alterations of *crk2* are caused by decreased phloem loading, which resulted in starch accumulation in source organs, and subsequent sink tissue starvation. Our results propose CRK2 as negative regulator of CALS1 and CALS3 regulating source to sink transport, which impacts plant growth and development.

## Introduction

Intercellular communication is a pivotal factor impacting development and stress responses of multicellular plants (Van Norman et al., 2011; Takahashi and Shinozaki, 2019; Tee and Faulkner, 2024). Generally, intercellular communication pathways can be divided into apoplastic, where molecules and their perception are located in the apoplastic space; and symplastic pathways, where molecules are transported between adjacent cells through cytoplasm. In plants, symplastic communication between cells is facilitated by microscopic intercellular channels called plasmodesmata (PD) (Bayer and Benitez-Alfonso, 2024). PD allow the symplastic transport of various micro-and macromolecules, including sugars, further allowing phloem loading, a process important for source to sink transport (Braun, 2022; Miras et al., 2022).

Symplastic communication can be enzymatically controlled by callose synthases (CALSs), also known as glucan synthase-like (GSL), which are responsible for the deposition of callose, (1→3)-β-d-glucan, in the neck region of PD. While the roles of various CALSs in specific biological processes have been described (Záveská Drábková and Honys, 2017; Ušák et al., 2023), less is known about their regulation at the molecular level (Dong et al., 2008; Faulkner et al., 2013; Cui and Lee, 2016; Záveská Drábková and Honys, 2017), especially their interactions with other proteins and formation of molecular complexes (Hong et al., 2001b). Analysis of callose deposition, combined with analyses of mutants and gene expression, provide valuable insights into the roles of CALSs in distinct biological processes. The positive correlation between increased callose, thus blocked symplastic transport, and starch levels can serve as an example (Julius et al., 2018; Li et al., 2024; Tee et al., 2024) and underlines the role of CALSs in the carbohydrate partitioning (Braun, 2022; Miras et al., 2022).

Several receptor-like kinases (RLKs) and receptor-like proteins (RLPs) localize to PD (Fernandez-Calvino et al., 2011) or to close proximity of PD-localised proteins (Li et al., 2024). Some of them have been described to be involved in the regulation of PD-mediated transport (Faulkner et al., 2013; Grison et al., 2019; Hunter et al., 2019; Cheval et al., 2020), for example PD-located proteins (PDLPs) (Tee et al., 2023; Li et al., 2024), or the closely related CYSTEINE-RICH RECEPTOR-LIKE PROTEIN KINASE 2 (CRK2) (Hunter et al., 2019; Vaattovaara et al., 2019; Kimura et al., 2020), also known as altered seed germination 6 (Bassel et al., 2011). CRKs are one of the largest Arabidopsis RLK families (Shiu and Bleecker, 2003), whose members are involved in various stress and developmental processes (Bourdais et al., 2015; Zeiner et al., 2023; Zhang et al., 2023; Krasensky-Wrzaczek and Wrzaczek, 2024). As the majority of CRKs are predicted to contain an active protein kinase domain (Vaattovaara et al., 2019), recruitment and phosphorylation of interacting proteins have been proposed as their main mode of action. This can be demonstrated for CRK2, which has been shown to regulate the production of apoplastic reactive oxygen species via phosphorylation of the NADPH oxidase RESPIRATORY BURST OXIDASE HOMOLOG D (Kimura et al., 2020). Furthermore, CRK2 interacts with several CALSs *in vivo* (Hunter et al., 2019), and phosphorylates CALS1 *in vitro* (Hunter et al., 2019). Importantly, the *crk2* mutant exhibits early flowering (Colina et al., 2026) and small rosette size phenotype (Bourdais et al., 2015; Kimura et al., 2020). The later one is also characteristic for the plants overexpressing *PDLP5* and *PDLP6* (Li et al., 2024). Additionally, members of the PDLP protein family have been previously linked to the regulation of CALS1 (Cui and Lee, 2016; Tee et al., 2023). The presence of CRK2 at or in close proximity to PD is not constitutive (Fernandez-Calvino et al., 2011; Hunter et al., 2019). Spatiotemporal analyses showed rapid stress-triggered enrichment of CRK2 at PD (Hunter et al., 2019). Accordingly, CRK2 has been proposed to participate in the regulation of stress-triggered callose deposition (Hunter et al., 2019; Kimura et al., 2020), yet its importance in plant growth and development has so far not been thoroughly investigated.

In this study, we investigated the genetic interaction between CRK2 and the phylogenetically closely related CALS1 (GSL6) and CALS3 (GSL12). These CALSs were previously identified as proteins interacting with CRK2 *in vivo* (Hunter et al., 2019) and we found both to be phosphorylated by CRK2 *in vitro*. Additionally, *CALS1* and *CALS3* shared overlapping expression patterns, predominantly localising the activities of their promoters into plant vasculature. We also demonstrated that while callose levels in *crk2* epidermal cells were elevated, callose levels in the *crk2 cals1.5* double mutant were similar to the levels in wild type plants, supporting a role for CRK2 as a negative regulator of callose deposition, as supported by the analysis of enzymatic activity in the heterologous yeast system. Furthermore, independent introduction of *cals1* or *cals3* into the *crk2* genetic background resulted in reversion of the small rosette size, reduced number of true leaves and short primary root, as well as the starch accumulation phenotype of *crk2*, which we describe here. Importantly, vasculature loading was noticeably decreased in *crk2*, suggesting decreased source-sink transport. This observation was in line with the negative impact of *crk2* shoot on wild type root growth in our grafting experiment. Our data show that CRK2, together with CALS1 and CALS3, controls symplastic loading of vasculature, distribution of growth and developmental signal molecules, especially sugars, and thereby impacts plant ontogenesis, as demonstrated by the reversion of *crk2* phenotypes.

## Results

### CRK2 physically interacts with and phosphorylates CALS1 and CALS3 in vitro

In our previous immunoprecipitation coupled with mass spectrometry analyses, we identified CALS1, CALS3 and CALS12 as *in vivo* interaction partners of CRK2 (Hunter et al., 2019). As the regulation of CALS1 was proposed previously (Hunter et al., 2019), we investigated whether CRK2 can potentially also regulate CALS3 and CALS12. Firstly, we carried out phylogenetic analyses of CALS protein sequences to explore their evolutionary divergence.

The CALS protein family in Arabidopsis consists of twelve members forming three distinct subclades (Figure 1A) in a phylogenetic maximum likelihood tree. CALS1 and CALS3 were identified to be members of a single subclade (Figure 1A), while CALS12 was placed in a different subclade. Since the likelihood for the conservation of regulatory elements is higher between closely related proteins, we subsequently focused on CALS1 and CALS3.

**Figure 1:**
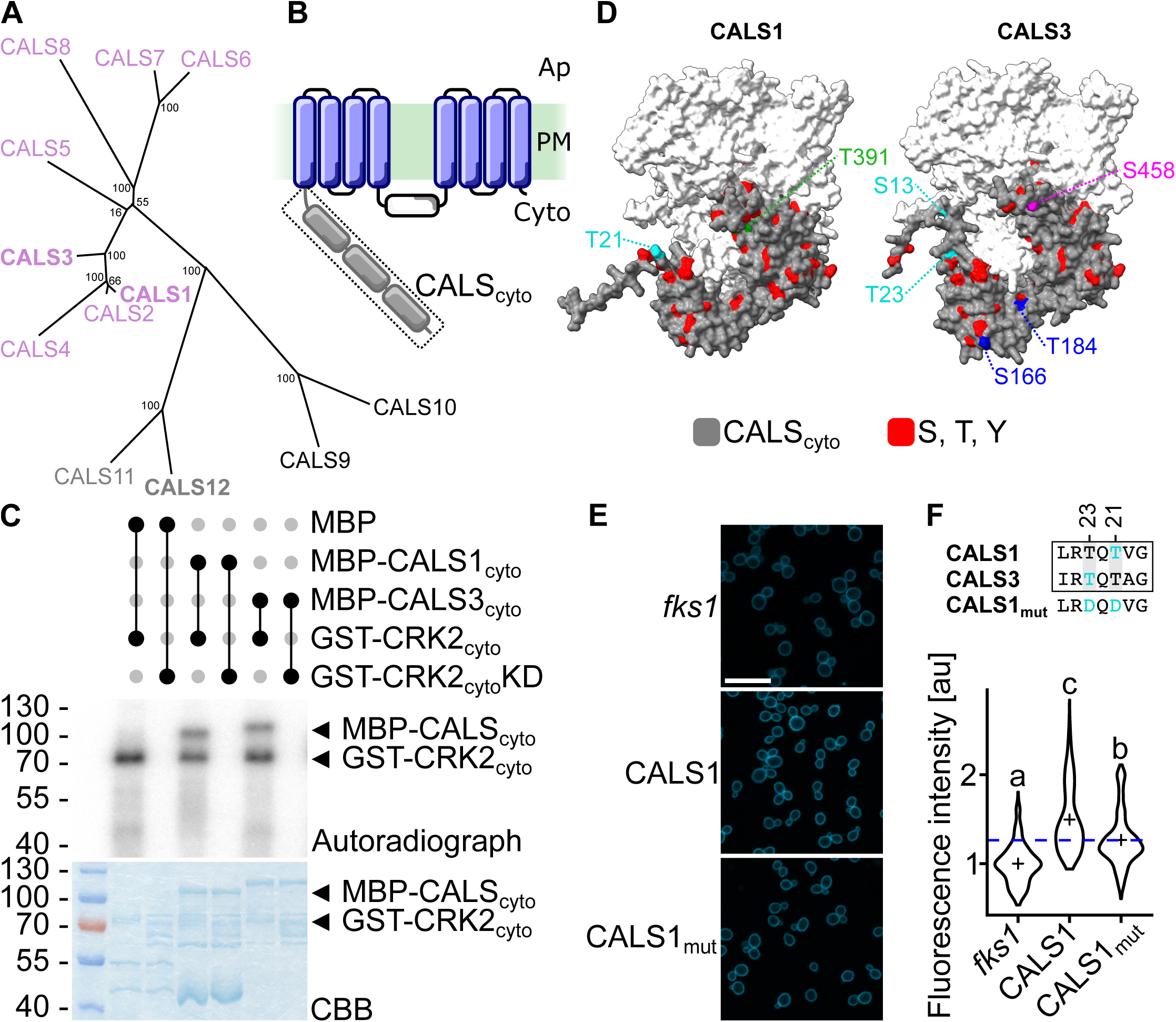
CRK2 phosphorylated CALS1 and CALS3 *in vitro*. (A) Phylogenetic analysis of the CALS family based on the protein sequences. Previously identified CALS1, CALS3 and CALS12 as *in vivo* interaction partners (Hunter et al., 2019) are indicated in bold. Different colours indicate distinct subclades. Numbers at the nodes show bootstrap values. (B) Cytoplasmatic (cyto) N-terminal region, containing 3 annotated domains (grey). This region is followed by the transmembrane domains (blue) anchoring CALS in plasma membrane (PM), cytoplasmic loop, another region containing transmembrane domains, and C-terminal cytoplasmic tail. Individual transmembrane domains are in cytoplasmic and apoplastic (Ap) regions connected by short linkers. (C) *In vitro* kinase assay of cytoplasmic region of CRK2 containing active kinase domain (CRK2_cyto_), kinase dead CRK2 (CRK2_cyto_KD), and N-terminal region (indicated in panel B) of CALS1 (CALS1_cyto_) and CALS3 (CALS3_cyto_). CBB – Coomassie brilliant blue, MBP – maltose binding protein, GST – glutathione S-transferase, relative molecular weight is in kDa. (D) Availability of CALS1 and CALS3 phospho-sites (red) in N-terminal region (grey) of indicated predicted structures of CALSs. Identified phospho-sites (this study) are highlighted. (E) Aniline blue staining (blue) of glucan in *fks1* mutant yeast strain complemented with CALS1_WT_ and CALS1_mut_. Bar – 20 µm. (F) Quantification of aniline blue fluorescence in yeasts. CALS1 and CALS3 identified conserved sites and altered sequence in CALS1_mut_ are in upper panel. Number indicates position in the sequence of respective CALS. Kruskal–Wallis test (Df = 2, H = 384.48) with Wilcoxon test. Cross indicates mean, blue dashed line indicates mean value of CALS1_mut_, different letters indicate statistical significance p < 0.05, n ≥ 250.

CRK2 has previously been found to phosphorylate the N-terminal cytoplasmic region of CALS1 *in vitro* (Hunter et al., 2019). Therefore, we focused on the cytoplasmic N-terminal regions (Figure 1B) of CALS1 and CALS3 in detail. We compared the amino acid sequences of CALS1 and CALS3 (Figure S1A), and indicated annotated domains (Figure 1B), based on the protein sequence alignment used for the construction of the phylogeny (alignment containing all Arabidopsis CALS is available as Data S1). CALS1 and CALS3 contained different numbers of potential phosphorylation sites (Figure S1A) and shared ∼85 % potentially conserved phosphorylation sites. Notably, both CALS1 and CALS3 have been identified as *in vivo* phosphoproteins (Umezawa et al., 2013; Wang et al., 2013) and are listed in the publicly available database PhosPhAt 4.0 (Zulawski et al., 2013) (Figure S1A). Additionally, we used structural predictions (Jumper et al., 2021), and highlighted the potential phospho-sites in N-terminal regions of CALS1 and CALS3 (Figure 1D, Figure S1B). We observed surface exposure of potential phosphorylation sites. Thus, our analysis pointed towards phosphorylation as an intersecting regulatory mechanism of CALSs.

To determine whether CRK2 could directly phosphorylate CALS3, similarly to CALS1 (Hunter et al., 2019), we performed *in vitro* kinase assays. The C-terminal cytoplasmic region of CRK2 harbouring the protein kinase domain (CRK2_cyto_), but not an inactive variant (CRK2_cyto_KD), was capable of phosphorylating the N-terminal cytoplasmic region of CALS3 (CALS3_cyto_) *in vitro*, similar to phosphorylation of CALS1_cyto_ (Figure 1C). To test the promiscuity of CALS1 and CALS3 as kinase substrate (Figure S1C), we used the cytoplasmic region of the RLK FERONIA (FER_cyto_). Since we observed lower activity for FER compared to CRK2 *in vitro* (Figure S1C, left panel), we separately exposed the gels with CRK2 (Figure S1C, middle panel) and FER (Figure S1C, right panel). Unlike for CRK2, we did not observe *in vitro* transphosphorylation of CALS1 or CALS3 by FER, supporting the specificity of CRK2 mediated regulation.

Additionally, we identified CRK2-dependent *in vitro* phosphorylation sites by phosphoproteomics (Figure S1A, highlighted by grey asterisks; Data S2). In particular phosphorylation sites located in the Vta1 (VPS20-associated protein 1) and FKS1 (1,3-beta-glucan synthase domain) domains might impact protein complex formation or callose synthase activity. Interestingly, one of the *in vitro* CRK2-dependent sites we identified, threonine 23 (T23) of CALS3, was previously described as an *in vivo* phosphorylation site (Wang et al., 2013).

To test the biological significance of identified CRK2 targeted sites, we adopted yeast heterologous system that employs *fks1* mutant yeasts, a strain deficient in β-1,3-glucan synthase with over all decreased glycan content (Inoue et al., 1995), and complementation with Arabidopsis CALS1 (Figure 1E). We selected tested sites identified in our phosphoproteomics analyses based on (i) availability to the intermolecular interaction and possible transphosphorylation (Figure 1D, Figure S1B), (ii) conservation of those sites between CALS1 and CALS3 (Figure 1F, upper panel; Figure S1D), and (iii) *in planta* identification of phosphorylation (Figure S1A). Based on these criteria, we selected CALS1 T19 and T21 (Figure 1F, upper panel). Those sites are not in CALS12 (Figure S1D). We used site-directed mutagenesis to generate CALS1, in which those amino acids were changed for phospho-mimicking aspartic acid (D) and designed it as CALS1_mut_ (Figure 1F, upper panel). After aniline blue staining of glucan of complemented yeasts with indicated construct (Figure 1E), we observed an increase of fluorescent signal in yeast complemented with both CALS1_WT_ and CALS1_mut_, compared to the *fks1* background (Figure 1F, bottom panel). Lower fluorescence of CALS1_mut_, reflecting the lower glucan content, suggested a negative regulatory role of the CALS1 T19 and T21 phosphorylation. Observed differences were not caused by the different expression levels, as confirmed by immunoblotting(Figure S1E).

Taken together, our analyses suggest that CALS1 and CALS3 could be directly phosphorylated, and thus regulated, by CRK2.

CALS1 *and* CALS3 *expression is not affected by CRK2* While our results suggested that CRK2 could regulate CALS1 and CALS3 directly, we also investigated whether *CALS1* and *CALS3* transcript abundance was altered in *crk2* compared to wild type.

We observed similar levels of *CALS1* and *CALS3* transcript levels between wild type and *crk2* (Figure 2A). This suggested, that CRK2 was not involved in controlling *CALSs* transcript abundance. Furthermore, we observed comparable transcript abundance of *CALS1* in wild type and *cals3.1*, and, similarly, *CALS3* transcript abundance in wild type was similar to *cals1.5*. This suggested, that the loss of one *CALS* gene is not compensated for by the upregulation of the expression of the other *CALS* gene.

**Figure 2:**
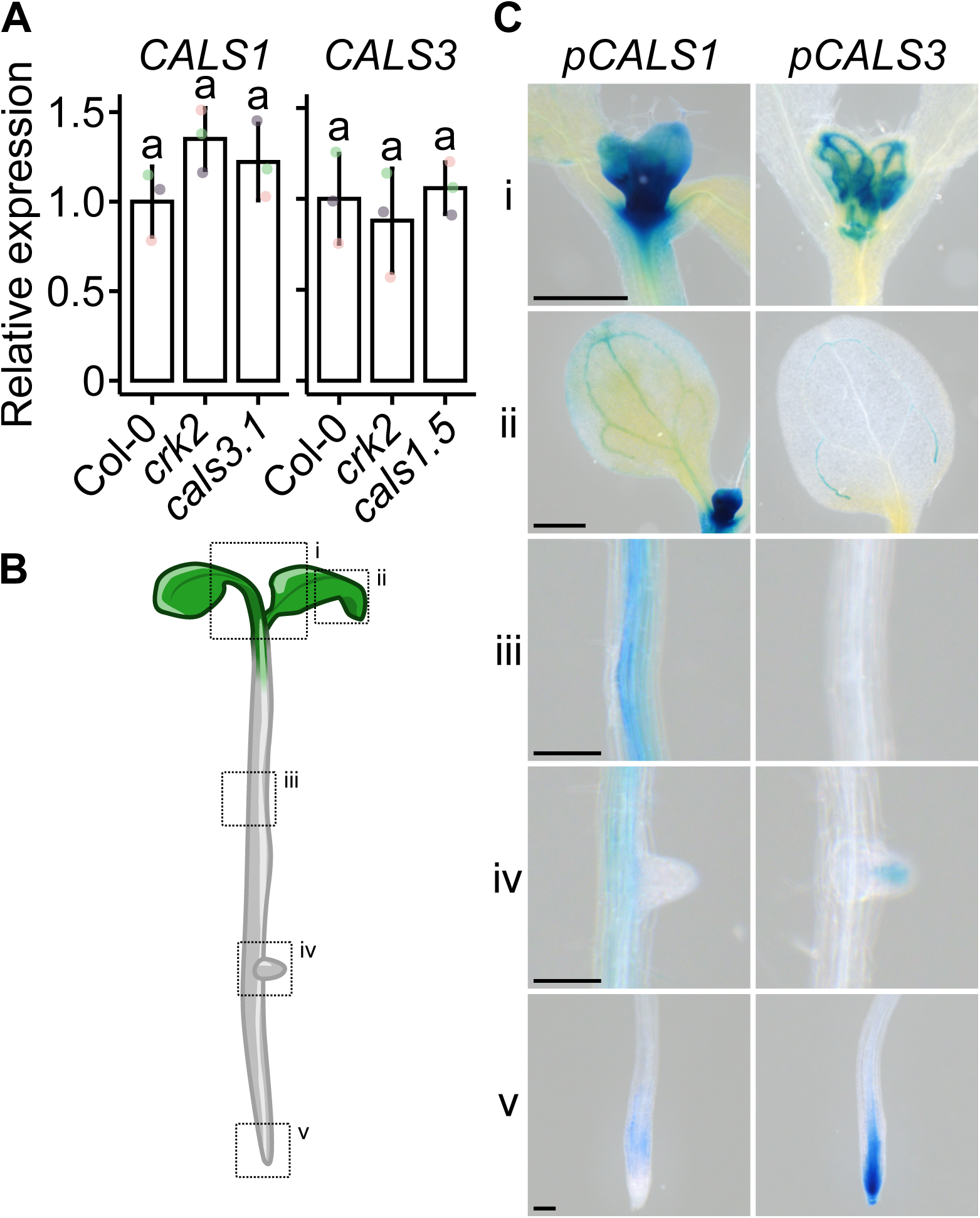
Expression levels of *CALS1* and *CALS3*, *CALS* with overlapping expression pattern, were not altered by CRK2. (A) Analysis of relative transcript abundance (relative to Col-0) between wild type and indicated mutants. Different colours indicate independent experiments, top of the bar indicates mean, whiskers show standard error. One-way ANOVA (Df = 2, 6; F_CALS1_ = 2.471, F_CALS3_ = 0.454) with Tukey’s HSD. Different letters indicate statistical significance p < 0.05, n = 3. (B) Schematic indication (i–v) of plant tissues exhibiting GUS staining, as analysed in the following panel. (C) Histological GUS staining analysis of *CALS1* and *CALS3* promoter driven expression of GUS reporter in shoot apieces, including emerging first true leaves and their vasculature (i), cotyledons (ii), primary root (iii), emerging lateral roots (iv) and primary root tip (v). (i, ii) Bar – 0.5 mm. (iii, iv, v) Bar – 0.1 mm.

To gain more comprehensive insight into *CALS1* and *CALS3* gene expression, we analysed their spatial expression under standard growth conditions. We generated transgenic plant lines expressing the β-glucuronidase (GUS) marker gene under the control of the native *CALS* promoters, *pCALS1* or *pCALS3*, together with the 3’UTR region of the respective gene. As indicated in schematic representation (Figure 2B), we observed strong histochemical staining of GUS driven by the *CALSs* promoters in shoot apices (Figure 2C, panel i), especially in vasculature of emerging first true leaves, and vasculature of cotyledons (Figure 2C, panel ii). Interestingly, *CALS1* promoter driven GUS expression not just in the vasculature, but widely throughout the whole cotyledon. In the root (Figure 2C, panels iii and iv), promoter of *CALS1* showed predominant marker expression in the vasculature, while *CALS3* promoter activity was detected mainly in the tips of emerging lateral roots (Figure 2C, panel iv). In the root tip, a complementary expression pattern was observed, where the expression of *CALS1* promoter was located in the elongation zone, in contrast to the *CALS3* observed in the meristematic zone (Figure 2C, panel v). Additionally, *CALS1* promoter showed higher GUS expression levels compared to *CALS3* promoter, reflected by the generally more pronounced histological staining between respective transgenic lines (Figure 2C).

Our analysis showed a partially overlapping expression pattern of *CALS1* and *CALS3*, especially in the vasculature, suggested their comparable or complementary functions in similar biological processes.

### CRK2 *and* CALS1 *modulate symplastic connectivity*

To analyse the relationship between CRK2 and selected CALSs, we used a genetics approach and generated double mutants *crk2 cals1.5* and *crk2 cals3.1* (Figure S2A). As the main role of CALSs resides in the regulation of symplastic communication through callose deposition at PD, we analysed callose accumulation and intercellular transport.

First, we determined basal callose levels in epidermal cells of the cotyledons of 7-day-old seedlings as an approximation of the number of aniline blue-stained spots (Figure 3A), as described previously (Hunter et al., 2019). We did not observe any significant differences in callose levels between *cals* mutants and wild type. However, *crk2* and the *crk2 cals3.1* double mutant displayed higher levels of callose compared to the wild type (Figure 3B). By contrast, callose levels were decreased in *crk2 cals1.5* compared to *crk2*, to a wild type level, suggesting that the increased level of callose in epidermal cells of *crk2* was caused by a lack of negative regulation of CALS1. A similar trend, based on the intensity of detected deposits (see Materials and Methods for details) was observed also in the mesophyll (Figure S2B, C). Interestingly, we did not observe any statistically significant differences between the tested genotypes for aniline blue staining in the vasculature associated callose (Figure S2B, D).

**Figure 3:**
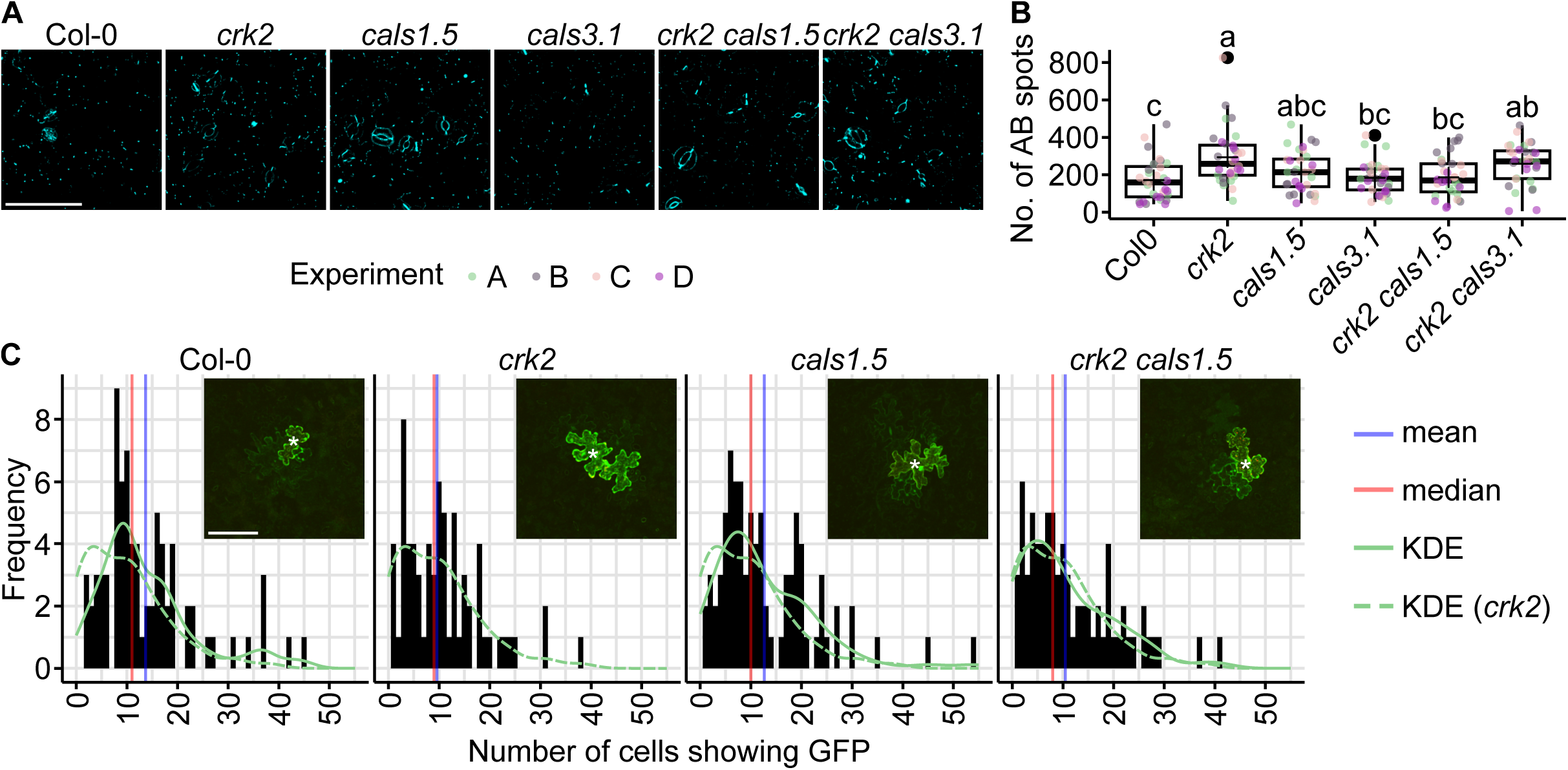
Increased callose deposition in epidermal cells of *crk2* did not significantly affect symplastic communication. (A) Aniline blue staining (blue) of callose deposits. (B) Quantification of aniline blue staining. One-way ANOVA (Df = 5, 185; F = 5.815) with Tukey’s HSD. Different letters indicate statistical significance p < 0.05, n = 32. Different colours indicate independent experiments, horizontal line indicates median, cross indicates mean, whiskers show minimum and maximum value, black dots indicate outliers, box indicates upper and lower quartiles. (C) Symplastic movement of GFP (green) in epidermal cells. Transformed cells are marked by the endoplasmic reticulum-targeted RFP (red) and indicated by asterisks (*). Bootstrap analyses of median where individual genotypes were compared to *crk2* show no statistical significance (not shown in respective panel), n ≥ 77. KDE – Kernel Density Estimation. (A, C) Bar – 100 µm.

To test whether symplastic transport was altered by CRK2 via CALS1, we used particle bombardment of epidermal cells (Tee et al., 2022). This method can be used to investigate the capability of symplastic transport by detection of GFP in non-transformed epidermal cells (Figure 3C, micrographs). Detected GFP is produced following gold particle-mediated transformation of individual cells (Figure 3C, micrographs, marked by asterisk). Presence of GFP in other cells is then scored and used as indicator of symplastic flux (Figure 3C). We observed subtle differences in the frequencies of cells exhibiting GFP signal between *crk2* and wild type or *cals1.5* plants. Importantly, the intermediate phenotype of *crk2 cals1.5* between *crk2* and *cals1.5* was reflected by a decreased number of cells showing GFP. This was visible especially after curve smoothing using kernel density estimation (KDE), which we used for easier characterisation of distribution between datasets of *crk2* (Figure 3C, dashed green curve) and the other genotypes (Figure 3C, solid green curve). We observed a similar unimodal distribution for wild type, *cals1.5* and *crk2 cals1.5*, while a bimodal distribution for *crk2*. The changes in symplastic transport between epidermal cells were reflected in no observed statistical significances between analysed genotypes, when we used bootstrapping analysis of the median values.

Taken together, the higher callose accumulation in epidermal cells of *crk2* was linked to CALS1. However, despite the higher callose accumulation in *crk2*, we observed only subtle impact of CRK2 on symplastic transport in epidermal cells of terminally developed cotyledons of 7-days-old plants.

### CALS1 and CALS3 impact CRK2-mediated developmental and growth processes

To assess the result of the genetic interaction, we performed phenotypic characterisation of the double mutant lines. The *crk2* mutant has been found to display reduced growth (Bourdais et al., 2015; Kimura et al., 2020) and reduced flowering time (Colina et al., 2026). Additionally, short day (SD) (Bourdais et al., 2015; Kimura et al., 2020) and long day (LD) photoperiods (Bourdais et al., 2015; Hunter et al., 2019; Kimura et al., 2020; Colina et al., 2026) were used for the analysis of *crk2*. Thus, we decided to perform comprehensive phenotypic analyses and integrate different photoperiods to study the genetic interaction of *CRK2* with *CALS1* and *CALS3*.

The *crk2* mutant displayed reduced rosette size under LD conditions (Bourdais et al., 2015; Kimura et al., 2020). In contrast to *crk2*, *cals1.5* and *cals3.1* showed no alteration of rosette fresh weight of 21-days-old plants compared to the wild type plants (Figure 4A, B). Interestingly, the independent introduction of *cals1.5* and *cals3.1* T-DNA insertional alleles into the *crk2* genetic background reverted the small rosette phenotype of *crk2* (Figure 4A1, B). We observed a similar reversion of *crk2* phenotype for rosette area (Figure S3A) and dry weight (Figure S3B). A similar trend was observed for plants grown in SD photoperiod (Figure 4C, D; Figure S3C, D). Previously, we observed that the small growth phenotype of *crk2* was more pronounced in SD conditions, compared to LD (Figure 4A-D). To evaluate the impact of different photoperiods on *crk2*, we normalized the fresh weight of *crk2* to the average value of the respective repeat of experiment and light condition of wild type; and compared the obtained values for *crk2* between LD and SD (Figure 4E). Intriguingly, the shortened light phase noticeably pronounced the fresh weight phenotype of *crk2* (Figure 4E). Correspondingly, this was also observable for normalised rosette area (Figure S3E) as well as dry weight (Figure S3F).

**Figure 4:**
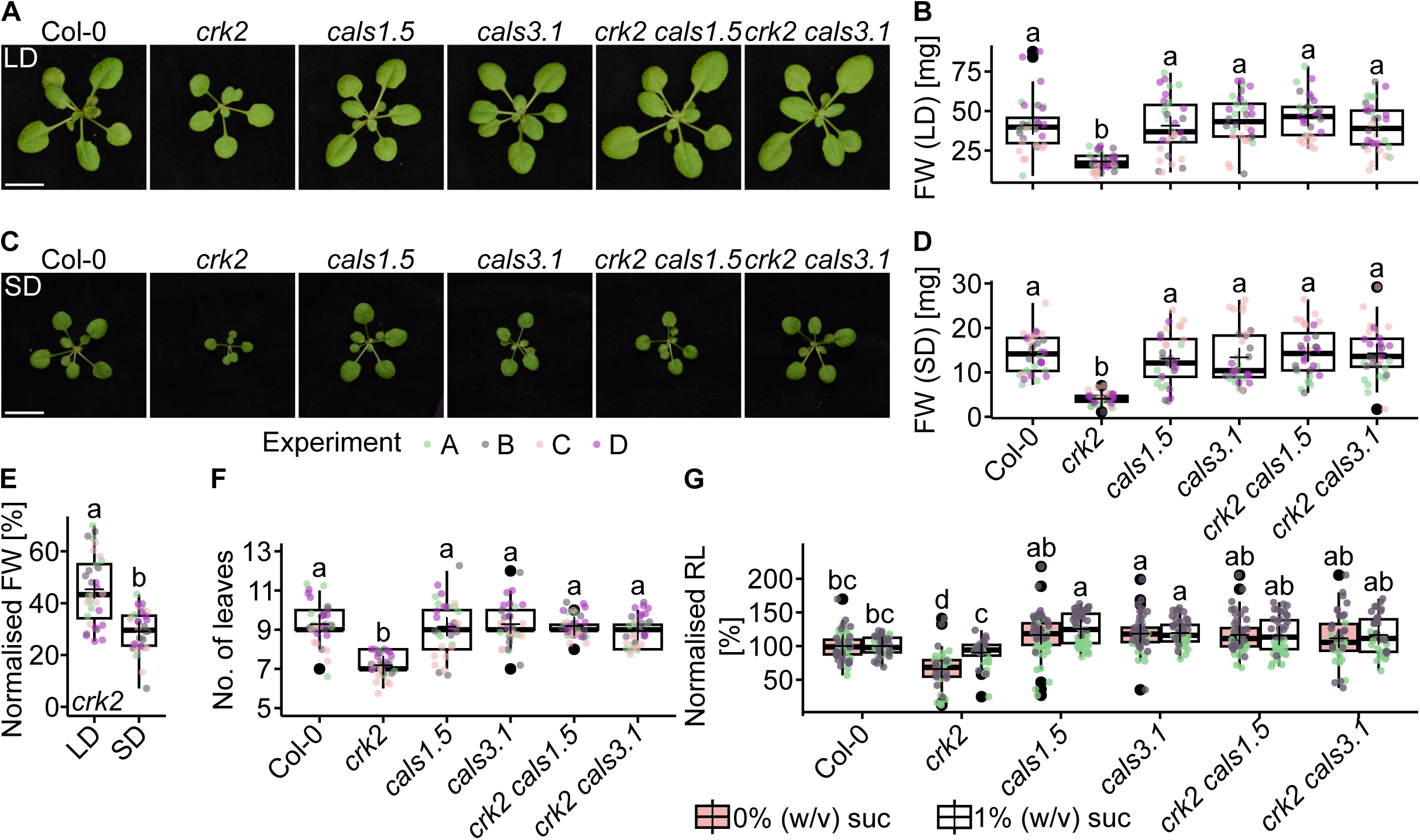
Introduction of *cals1.5* and *cals3.1* alleles into *crk2* genetic background reverted small rosette, developmental delay and shorter primary root phenotype of *crk2*. (A) Rosettes of 21-days-old plants grown in long day (LD) conditions. (B) Fresh weight (FW) of rosettes of 21-days-old plants grown in LD conditions. Kruskal–Wallis test (Df = 5, H = 63.079) with Wilcoxon test. (C) Rosettes of 21-days-old plants grown in short day (SD) conditions. (D) FW of rosettes of 21-days-old plants grown in SD conditions. (A, C) Bar – 10 mm. (D) Kruskal–Wallis test (Df = 5, H = 74.661) with Wilcoxon test. (E) Normalized FW of 21-days-old plants grown in indicated photoperiod. Wilcoxon test (W = 864). (F) Number of the leaves of 21-days-old plants grown in LD conditions. Kruskal–Wallis test (Df = 5, H = 71.326) with Wilcoxon test. (G) Normalized primary root length (RL). Two-Way ANOVA (treatment: Df = 1, 529, F = 6.330; genotype: Df = 5, 529, F = 33.319; interaction: Df = 5, 529, F = 2.745) with Tukey’s HSD, n ≥ 35. (B, D, E, F) n = 32. (E, G) Values normalized to the mean value of Col-0 of the respective repeat and treatment. (B, D, E, F, G) Different letters indicate statistical significance p < 0.05. Different colours indicate independent experiments, horizontal line indicates median, cross indicates mean, whiskers show minimum and maximum value, black dots indicate outliers, box indicates upper and lower quartiles.

The *crk2* mutant was recently described to have fewer leaves at flowering initiation compared to wild type (Colina et al., 2026). Thus, we decided to also analyse the number of the leaves of 21-days-old plants (Figure 4A). We observed a significantly decreased number of leaves in *crk2* compared to the other genotypes, including double mutants (Figure 4F). The decreased number of true leaves underlined the impact of the genetic interaction between CRK2 and CALSs on plant development, beyond simple modulation of growth.

Since *crk2* also displayed a shorter primary root compared to wild type in normal growth conditions (Hunter et al., 2019), we tested whether genetic interaction between *CRK2* and *CALSs* affected also this phenotype. We observed that in media without sucrose, *crk2* exhibited a significantly shorter primary root, compared to wild type (Figure 4G). Similar to rosette size, the introduction of *cals1.5* and *cals3.1* mutant alleles into *crk2* genetic background reverted the shorter primary root length phenotype of *crk2* (Figure 4G), further supporting interplay of *CRK2* and *CALS* in the same or overlapping developmental processes. Additionally, supplementation of media with an external carbon source in the form of 1% (w/v) sucrose (suc), recovered the observed *crk2* phenotype, while it had no impact on the double mutant and respective *cals* parental lines (Figure 4G). We did not observe any differences in shoot development between the tested genotypes (Figure S3G). Further, we tested the impact of sucrose supplementation on the phenotype of 21-days-old plants (Figure S3H). We observed a partial reversion of rosette area (Figure S3I) and leaf number (Figure S3J) in *crk2* and plants expressing an inactive variant of CRK2 (CRK2^D450N^, *pCRK2::CRK2^D450N^-mVenus*/*crk2*), compared to wild type and CRK2 complementation plants (CRK2, *pCRK2::CRK2-mVenus* #1-22/*crk2*). The reversion was not to the level of wild type plants, but distinguishable from *crk2* and plants expressing CRK2^D450N^ grown on media without sucrose. This points towards an alternation of growth by the carbon allocation.

Taken together, CALS1 and CALS3, impact developmental and growth processes which are also regulated by CRK2.

### CRK2 controls starch accumulation

The availability and distribution of nutrients play a crucial role in growth. Overexpression of *PDLP6*, which is closely related to *CRK2* (Zeiner et al., 2023), resulted in a small rosette size phenotype (Li et al., 2024), similar to *crk2* (Figure 4A-D). Since plants overexpressing *PDLP6* (Li et al., 2024), and plants expressing constitutively active CALS3 (Tee et al., 2024) showed on top of reduced growth also enhanced starch accumulation, we analysed the starch accumulation by histological Lugol’s staining and biochemical analyses.

In comparison to wild type, we observed pronounced starch accumulation in different developmental stages in *crk2*. Specifically, we observed starch buildup in the cotyledons of 7-days-old (Figure S4A) and 10-days-old *crk2* plants, compared to wild type (Figure 5A-D). Since we previously used sucrose-supplemented medium, we analysed starch accumulation in those conditions. We did not observe any significant impact of sucrose supplementation on the observed trend (Figure S4B), compared to media without sucrose supplementation (Figure 5A-D). Additionally, 28-days-old *crk2* plants showed similar trend (Figure S4C). Intriguingly, the observed starch accumulation in *crk2* was independent of diurnal starch metabolism dynamics, as plants collected at the end of the day (EoD, Figure 5A, B) and end of the night (EoN, Figure 5C, D) showed a similarly pronounced accumulation of starch compared to wild type and the other genotypes. Similarly to growth and developmental phenotypes, we observed the reversion of starch accumulation by the introduction of *cals1.5* and *cals3.1* mutant alleles into *crk2* mutant background (Figure 5A-D).

**Figure 5:**
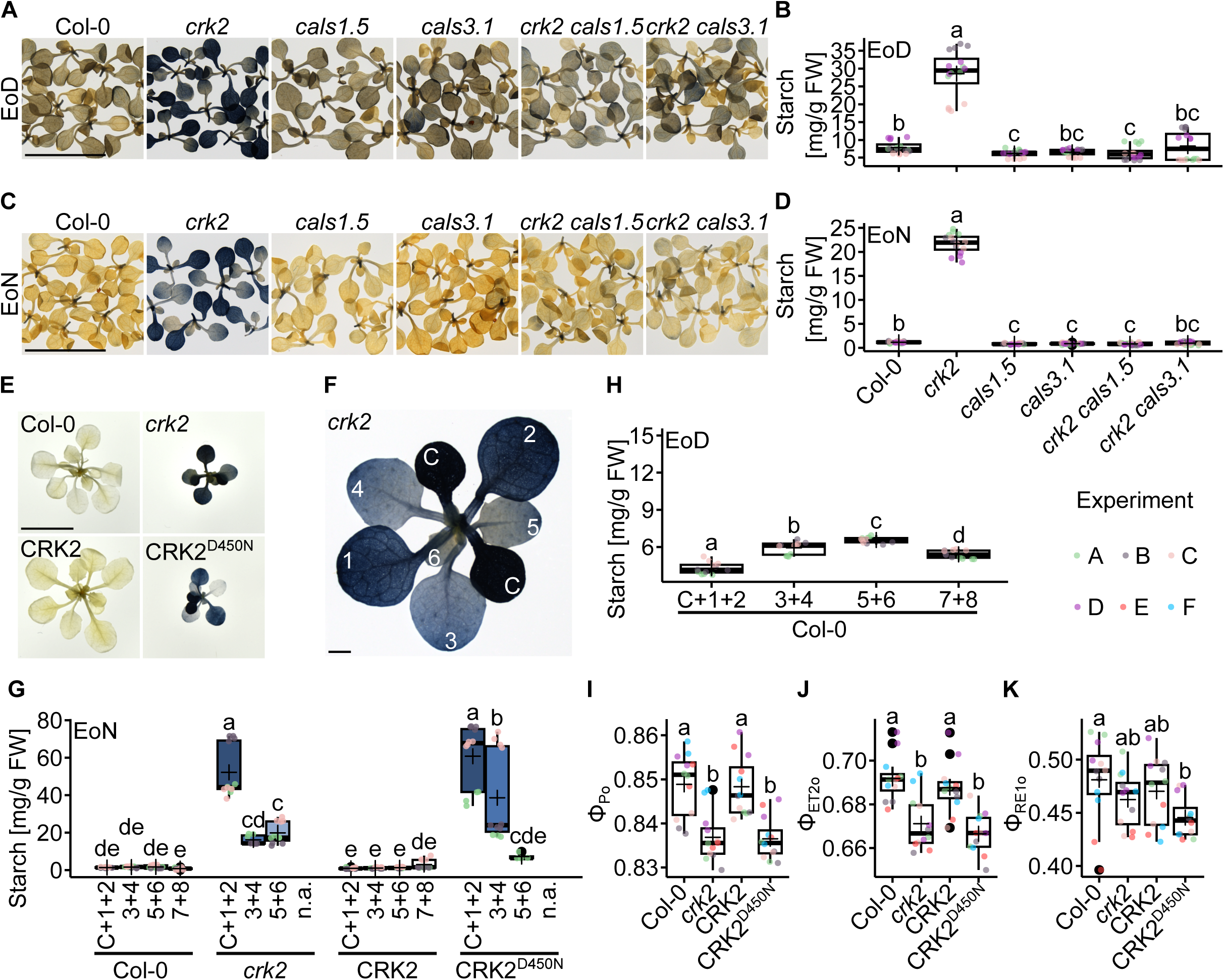
Ectopic starch accumulation of *crk2* was reverted by the independent crossing with *cals1.5* and *cals3.1* parental lines. (A) Lugol’s staining of 10-days-old plants at the end of the day (EoD). (B) Biochemical quantification of starch in 10-days-old plants at the end of the day. Kruskal–Wallis test (Df = 5, H = 45.3). (C) Lugol’s staining of 10-days-old plants at the end of the night (EoN). (D) Biochemical quantification of starch in 10-days-old plants at the end of the night. Kruskal–Wallis test (Df = 5, H = 51.487). (E) Lugol’s staining of 22-days-old plants expressing kinase active variant of CRK2 (CRK2, *pCRK2::CRK2-mVenus* #1-22/*crk2*), or (CRK2^D450N^, *pCRK2::CRK2^D450N^-mVenus*/*crk2*). (F) Lugol’s staining of 22-days-old *crk2* plant. (A, C, E, F) Bar – 10 mm. (G) Biochemical quantification of starch in indicated leaves of 22-days-old plants at the end of the night. The division of the samples by the certain leaves is indicated in panel F. Two-Way ANOVA (leaf: Df = 3, 138, F = 62.44; genotype: Df = 3, 138, F = 157.36; interaction: Df = 7, 138, F = 18.12) with Tukey’s HSD. (H) Biochemical quantification of starch in indicated leaves of 22-days-old wild type at the end of the day. One-way ANOVA (Df = 3, 44; F = 72.38) was followed by the Tukey’s HSD. (F, G, H) Number indicates the respective leaf. C – cotyledon. (I, J, K) – Photosynthesis efficiency parameters. (I) Maximum dark-adapted quantum yield of primary PSII photochemistry (Φ_Po_). Kruskal–Wallis test (Df = 3, H = 24.711). (J) Quantum yield of the electron transport flux from QA to QB. Kruskal–Wallis test (Df = 3, H = 24.702). (K) Electron transport to PSI. Kruskal–Wallis test (Df = 3, H = 7.8467). (B, D, I, J, K) Kruskal–Wallis test was followed by Wilcoxon test. (B, D, G, H, I, J, K) Different letters indicate statistical significance p < 0.05. Different colours indicate independent experiments, horizontal line indicates median, cross indicates mean, whiskers show minimum and maximum value, black dots indicate outliers, box indicates upper and lower quartiles, n ≥ 7.

Interestingly, we observed increased levels of starch in cotyledons of *crk2* mutants, compared to the developmentally younger true leaves (Figure 5A, C). This gradient was even more pronounced in older plants (Figure S4C). Additionally, we focused on the previously described CRK2 complementation lines (Hunter et al., 2019). Since we did not observe any significant diurnal effect on the phenotype observed for *crk2*, we analysed plants at the end of the night (Figure 5E-G). We did not observe an accumulation of starch in wild type and CRK2 complemented plants. By contrast, strong starch accumulation was observed in *crk2* and plants expressing an inactive variant of CRK2^D450N^. Further analysis revealed a decreasing starch accumulation gradient towards the developmentally younger leaves in *crk2* (Figure 5F, G). Interestingly, when we analysed the starch content of wild type plants at the end of the day, when we expected maximal starch accumulation, we observed an increasing character of starch accumulation trend (Figure 5H), in comparison to decreasing trend observed in *crk2* (Figure 5G). This suggest that the accumulation in *crk2* is linked to sugar distribution. This gradient was independent of sucrose supplementation in media (Figure S4D).

As developmentally younger tissues of plants are sinks, we asked whether the observed effect was connected to (i) the age of the tissue or to (ii) source-to-sink transport. Thus, we analysed the root tips as a pure carbon sink tissue. We observed decreased levels of starch in *crk2* and CRK2^D450N^, compared to roots of wild type and CRK2 complemented plants (Figure S4E), stressing the role of the sugar transport over the age-dependent starch accumulation.

To further investigate the connection between increased levels of callose and starch, we applied salicylic acid (SA), which was previously described as a potent stimuli of callose deposition (Dong et al., 2008; Cui and Lee, 2016). Application of SA, but not its inactive isomer 4-hydroxybenzoic acid (4-OHBA), increased starch levels in wild type as expected (Figure S4F, G). Interestingly, starch levels in SA-treated wild type plants were still lower compared to *crk2*, for which we did not observe any significant impact of SA treatment (Figure S4G). Although we observed the expected increase of starch levels in wild type plants treated with SA, we did not observe statistically significantly elevated callose levels in epidermal cells of cotyledons (Figure S4H).

Increased starch levels could be caused by increased photosynthetic capacity. Alternatively, accumulation of saccharides can lead to the downregulation of photosynthesis. Therefore, we analysed photosynthetic parameters by fast chlorophyll fluorescence induction kinetics (OJIP) to characterise the efficiency of photosystem (PS) II and PSI, electron transport and the redox state of the photosystems. The *crk2* and CRK2^D450N^ lines exhibited lower maximum quantum yield of primary PSII photochemistry (Φ_Po_) (Figure 5I), and quantum yield of the electron transport flux from QA to QB (Φ_ET2o_) (Figure 5J), compared to wild type and CRK2 complemented plants. The effect on electron transport to PSI (Φ_RE1o_) was less pronounced (Figure 5K). In line with those changes, the photoprotection by non-photochemical quenching (NPQ) during light adaptation and relaxation was generally comparable between genotypes (Figure S5).

The decrease of photosynthetic efficiency was in line with the proposed predominant role of sugar transport in *crk2*, which caused the increased starch levels. Taken together, our analysis highlighted the role of kinase active CRK2 as an emerging central regulator of starch accumulation *via* controlling sugar distribution.

### CRK2 *impacts sink-source transport*

Symplastic phloem loading and the subsequent transport of nutrients from terminally developed source tissues could be a limiting factor for the development and growth of sink tissues. PD have central role in this process (Braun, 2022), thus we investigated the contribution of CRK2 in long distance transport. As we did not observe abnormal vein patterning in *crk2* cotyledons (Figure S6A), nor altered root architecture (Figure S6B), or altered meristem length (Figure S6C), we concluded that transport within vasculature, rather than its structure, could play a predominant role.

To test vascular transport efficiency, we adopted a modified version of the 5(6)-carboxyfluorescein diacetate (CFDA) assay (Kalmbach et al., 2023; Hajný et al., 2024), and applied CFDA directly into the exposed root vasculature (Figure 6A). This approach allowed us to bypass symplastic loading. After 2 hours following CFDA application, we observed entry of CFDA into the root vasculature, as fluorescence of carboxyfluorescein, and analysed fluorescence intensity in the root tip by laser scanning confocal microscopy (Figure 6B). We did not observe any significant differences in mobility of tracer within the vasculature or during post-vascular movement, as the fluorescent intensity (Figure 6C) and the distribution of signal (Figure 6B), respectively, in root tips were comparable between wild type and *crk2*.

**Figure 6:**
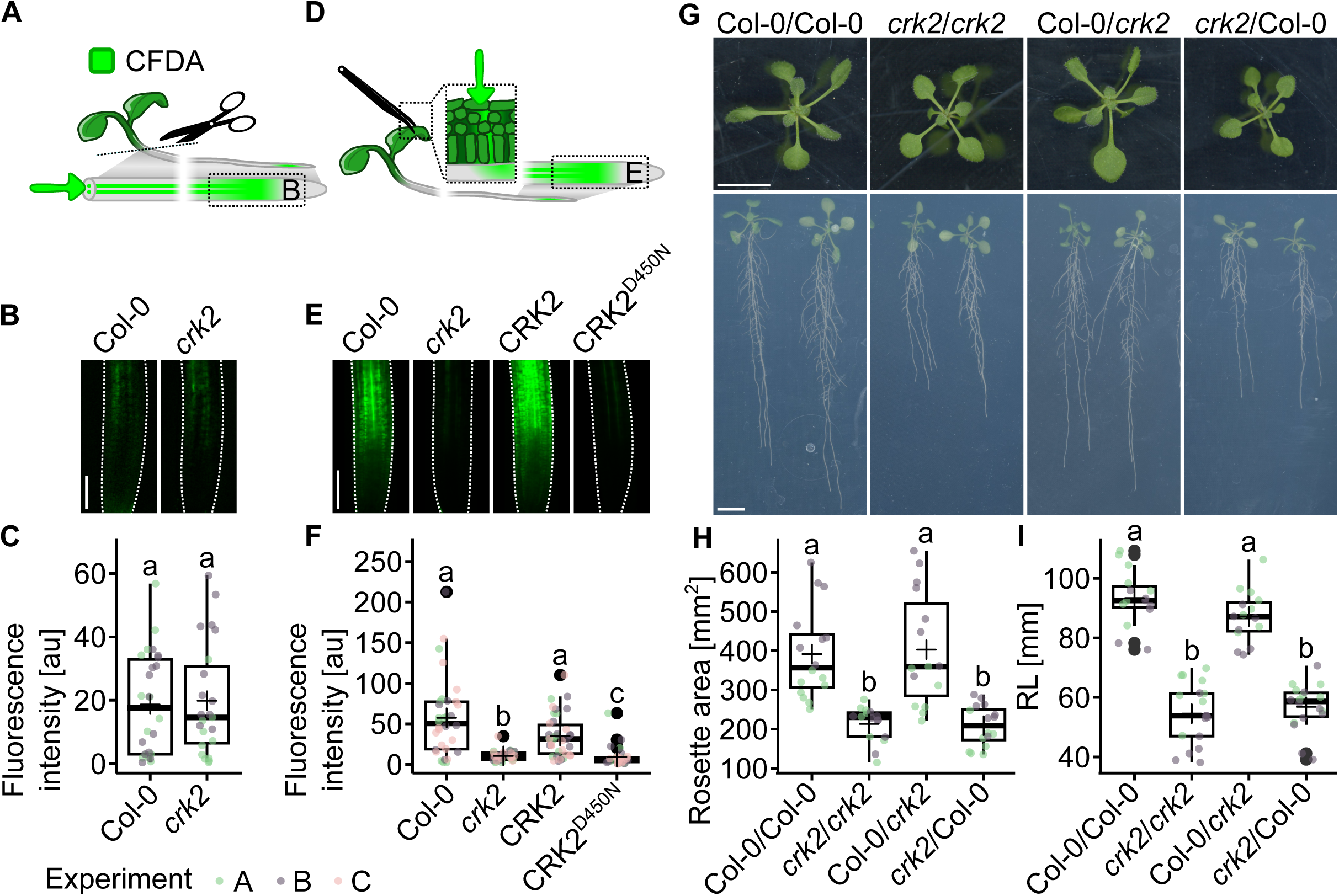
Long distance transport from cotyledons to root tip required kinase active CRK2. (A) Modified 5(6)-carboxyfluorescein diacetate (CFDA) assay, where CFDA was directly applied into root vasculature (indicated by the arrow), was used to analyse long distance transport. Fluorescence of applied tracer was subsequently analysed in the root tip (indicated in the box). (B) Fluorescence in the root tip after modified CFDA assay. (C) Quantification of fluorescence in the root tip after modified CFDA assay. Welch’s two sample t-test (Df = 44.817, t =-0.24785). (D) Classical CFDA assay uses tweezers to disrupt leaf tissue, which allows entry of dye. Contrary to the previously used approach (described in panel A), this protocol is testing symplastic loading of vasculature, which is subsequently used for the tracer transport. (E) Fluorescence in the root tip after CFDA assay in complementation lines with kinase active CRK2 (CRK2, *pCRK2::CRK2-mVenus* #1-22/*crk2*) and kinase inactive variant of CRK2 (CRK2^D450N^, *pCRK2::CRK2^D450N^-mVenus*/*crk2*). (B, E) Bar – 100 µm. (F) Quantification of fluorescence in the root tip of indicated genotypes after CFDA assay. Kruskal–Wallis test (Df = 3, H = 51.941) with Wilcoxon test. (G) Shoot (upper panel) and root (lower panel) of wild type plants and *crk2* after grafting. Bar – 10 mm. (H) Size of rosettes after grafting. One-way ANOVA (Df = 3, 62; F = 79.71) with Tukey’s HSD. (I) Primary root length (RL) after grafting. Kruskal–Wallis test (Df = 3, H = 26.108) with Wilcoxon test. (G, H, I) Labelled as shoot/root. (C, F, H, I) Different letters indicate statistical significance p < 0.05. Different colours indicate independent experiments, horizontal line indicates median, cross indicates mean, whiskers show minimum and maximum value, black dots indicate outliers, box indicates upper and lower quartiles, n ≥ 14.

Since we excluded the impact of long-distance transport within the vasculature, we applied CFDA onto cotyledons nicked by tweezers (Figure 6D). This approach requires symplastic loading. As described above (Figure 6B), the analysis was based on the quantification of the fluorescence intensity in the root tip (Figure 6E). Fluorescence intensities observed in the roots of *crk2* and *crk2* expressing the kinase inactive variant CRK2^D450N^ were significantly lower compared to wild type (Figure 6F). Expression of the kinase active variant of CRK2 in *crk2* background restored the transport capabilities to a levels comparable with wild type (Figure 6F). Furthermore, we tested the impact of the genetic interaction between *CRK2* and *CALSs* on vasculature loading (Figure S6D). We observed, that the independent introduction of *cals1* or *cals3* mutant alleles into *crk2* background increased the observed fluorescence intensities in the root tips to the level of *cals* parental lines compared to *crk2* (Figure S6E).

In addition, we tested the contribution of shoots and roots on the *crk2* phenotype (Figure 4) by grafting (Figure 6G-I). Grafting success was evaluated by analysing the phloem reconnection rate, which revealed no impact of *crk2* on this process (Figure S6F). Subsequently, we carried out reciprocal grafting between wild type and *crk2*, and analysed rosette area and primary root length. We observed reduced rosette size in *crk2* homografted plants, compared to wild type homografts consistent with previous experiments. Importantly, rosette size was not altered compared to wild type when wild type shoots were grafted on *crk2* roots (Figure 6G, upper part; Figure 6H). Similarly, we observed wild type-like root growth of *crk2* roots grafted to wild type shoots. However, when *crk2* shoots were grafted on wild type roots, we observed *crk2*-like rosette size (Figure 6G, upper part; Figure 6H) as well as a significant reduction of root growth, compared to wild type homografted plants (Figure 6G, lower part; Figure 6I).

These observations suggest CRK2 as a critical component controlling vascular loading in the source tissues, while long distance transport through the vasculature seemed to be unaffected in the *crk2* mutant.

## Discussion

CRK2, a member of one of the largest RLK subgroups (Zeiner et al., 2023), has previously been described to enrich at PD and regulate callose deposition following signal perception upon stress conditions (Hunter et al., 2019; Kimura et al., 2020). Importantly, the lateral distribution of CRK2 (Hunter et al., 2019), together with higher basal levels of PD-associated callose in *crk2* (Figure 3), highlighted the presence and functional importance of CRK2 in PD upon normal growth conditions (Figure 4). Interestingly, CRK5, CRK22 (Zhao et al., 2022), and CRK36 (Yeh et al., 2015; Lee et al., 2017) have been also reported to modulate PD functionality by the regulation of callose deposition upon stress conditions. However, the role of CRK2 in the regulation of plant growth and development by the modulation of callose deposition appears to be unique so far. While many studies on the characterisation of molecular mechanisms regulating callose deposition are focused on stress responses (Cui and Lee, 2016; Grison et al., 2019; Cheval et al., 2020), developmental aspects have been investigated to a lesser extent (Grison et al., 2019; Li et al., 2024). In this study, we focused on the characterisation of the genetic interaction between *CRK2* and two closely related *CALS1* and *CALS3* genes encoding CALSs previously identified as *in vivo* interaction partners of CRK2 (Hunter et al., 2019). Here, we show that both CALS were phosphorylated by CRK2 *in vitro* and impacted CRK2-mediated processes. *CALS1* is transcriptionally responsive to SA (Dong et al., 2008; Cui and Lee, 2016), and, together with *CALS3*, is one of the predominantly expressed *CALS* genes in Arabidopsis (Dong et al., 2008). CALS1 participates in the regulation of PD permeability in response to SA (Cui and Lee, 2016; Tee et al., 2023) and *Pseudomonas* treatment (Cui and Lee, 2016); and was further described to be involved in cytokinesis (Hong et al., 2001a; Hong et al., 2001b) and flowering time regulation (Murata et al., 2025). CALS3 has predominantly been described as an important regulator of plant growth via modulation of PD (Vatén et al., 2011; Saatian et al., 2023), however, mostly by using gain-of-function alleles.

Despite proposed functional specificities of respective genes, both *cals1* and *cals3* could rescue several *crk2*-related phenotypes. The previously described small rosette (Kimura et al., 2020), decreased number of leaves (Colina et al., 2026), and shorter primary root (Hunter et al., 2019) phenotypes of *crk2* were reverted by the independent introduction of *CALS1* or *CALS3* T-DNA insertion alleles into *crk2* mutant background (Figure 4). Noteworthily, introduction of *cals1* or *cals8* mutant alleles into plants overexpressing *PDLP5* rescued the PD permeability phenotype resulting from *PDLP5* overexpression (Cui and Lee, 2016). Similarly, independently silenced *CALS9* or *CALS10* in *cals12* resulted in a dwarf phenotype (Töller et al., 2008). This could propose the role of protein-protein interaction and heteromerization, which can modulate the activity of CALSs, as proposed previously (Töller et al., 2008). In contrast to *cals3.1*, we observed that the introduction of *cals1.5* into *crk2* mutant background decreased callose levels in *crk2 cals1.5* double mutant plants (Figure 3, Figure S2). This suggests that CALS1, rather than CALS3, could be regulated by CRK2. Consequently, we speculate that the reversion of callose phenotype in *crk2 cals1.3* was rather due to the lack of *CALS3* expression (Figure 2C, panel ii), and thus predominant role of CALS1 in epidermal and mesophyll cells, where the analysis was performed. Although we did not observe any difference in callose associated with vasculature, we were not able to test the specific cell interfaces. This analysis would provide unique insights into CRK2-mediated regulation of callose deposition. Collectively, this suggestsCALS1 and CALS3, together with their upstream regulators, as important regulators of plant growth and development. Further, it proposes CRK2 as negative regulator of callose deposition mediated by the regulation CALS1 and CALS3 and emphasized their participation in growth and developmental processes. CRK2 as the negative regulator of the callose deposition was further supported by the analysis of selected identified CRK2-dependent phospho-sites in a heterologous system (Figure 1E, F). Interestingly, the broader expression of *CALS1*, in particular in whole cotyledon, compared to the more narrow expression pattern of *CALS3* in the vasculature, suggested its importance for vasculature associated processes (Figure 2). We observed an increased callose deposition in *crk2* and its reversion only in *crk2 cals1.5* double mutant, pointing towards alterations of symplastic transport in those mutant lines. Surprisingly, symplastic movement of free GFP between epidermal cells was not significantly affected in *crk2* (Figure 3). While observed differences among the lines were not statistically significant, obvious trends could be noticed. Wild type, *cals1.5*, and *crk2 cals1.5* exhibited a unimodal, while *crk2* showed a bimodal character of distribution in histograms of GFP spread between cells, supporting the importance of those components in symplastic transport. Furthermore, it has to be noted that the increased callose levels in *crk2* compared to wild type were rather subtle in this dataset (Figure 3B), potentially attenuating the difference in GFP spread. This is in line with previous reports, where higher differences in callose levels were reflected in significant changes of the GFP spread (Xu et al., 2017; Cheval et al., 2020; Li et al., 2025), thus highlighting technical limitations in the analysis of symplastic permeability (Tee et al., 2022).

Plants accumulating callose, especially in close proximity to vasculature, have been shown to exhibit elevated levels of starch (Julius et al., 2018; Li et al., 2024; Tee et al., 2024), potentially as a result of decreased symplastic transport. In the light of independent studies, which showed involvement of various proteins in the simultaneous modulation of PD and carbohydrate metabolism (Lucas et al., 1993; Hofius et al., 2001; Julius et al., 2018; Tran et al., 2019; Li et al., 2024; Tee et al., 2024), we concluded that the excessive starch accumulation in *crk2* (Figure 5) could be a secondary effect of reduced PD-mediated transport (Figure 3). This was supported by a pharmacological approach, where application of SA, which has been shown to induce callose deposition and thereby modulate PD permeability (Dong et al., 2008; Cui and Lee, 2016), resulted in a significant increase of starch levels in wild type plants (Figure S4F, G). The pronounced accumulation of starch in *crk2,* compared to wild type plants and *cals* single mutants, was reverted in the double mutants and in lines complemented with kinase active CRK2. This phenomenon was observed in all tested developmental stages. Importantly, the enhanced starch accumulation in *crk2* was not caused by enhanced photosynthetic capacity (Figure 5I, J, K; Figure S5), but rather by defects in sugar allocation. Photochemical quantum yields decrease, and up-regulation of non-photochemical exciton dissipation, could be the result of feedback regulation in response to the increased sugar levels in *crk2*. Sugars, which would be used for growth and development (Figure 4), accumulated in source tissue and were allocated into starch (Figure 5). This can be supported by the described enhanced starch accumulation upon external carbon supplementation in the form of sucrose (Figure 4SD) (Laloum et al., 2022). Thus, analysis of sugar signaling pathways (Fichtner and Lunn, 2021) and proteins involved in sugar metabolism (Streb and Zeeman, 2012) would further provide valuable insight into alternations of signaling networks in *crk2*.

One of the examples of starch accumulating genotypes could be plants overexpressing *PDLP6* (Li et al., 2024), a gene encoding a protein closely related to CRK2 (Zeiner et al., 2023). *PDLP6* was found to be predominantly present in the vasculature, and its overexpression resulted in increased levels of vasculature-associated callose. Interestingly, starch accumulation was not uniform in *PDLP6* overexpression lines, but rather different between individual organs (Li et al., 2024). Compared to younger leaves, we observed increased starch accumulation in developmentally older leaves in *crk2* (Figure 5). In comparison to wild type, *crk2* plants exhibited nutrient accumulation in the form of starch in source tissues, which could lead to the sugar shortage in sinks. This was supported by the decreased starch levels in the root tips, a tissue solely dependent on sugar transport from source (Figure S4). *CALS7* is expressed in vasculature (Barratt et al., 2011; Xie et al., 2011) and *cals7* exhibited abnormal vascular structure and manifested reduced growth (Barratt et al., 2011; Xie et al., 2011) possibly as a result of tissue starvation (Barratt et al., 2011). Manifesting a similar phenotype, starvation could be a possible explanation also for *crk2* growth phenotype, as supplementation with external carbon source in the form of sucrose in the media (Figure 4G, Figure S3G-J) and extended day length (Figure 4E) reverted the *crk2* phenotype.

Constitutively active CALS3 affects shoot (Linh and Scarpella, 2022) and root (Vatén et al., 2011) vasculature, and as a consequence primary root length (Vatén et al., 2011). Importantly, *crk2* showed no evidence of abnormal vascular architecture (Figure S6) or transport from shoot to root within the vasculature (Figure 6C), as suggested by CFDA assay and transport of tracer from shoot to root after direct loading into the vasculature. Thus, we concluded that vascular transport *per se* was not causing developmental phenotypes of *crk2*. When we applied CFDA to source leaves, and its distribution therefore dependent on transport towards the vasculature and proper loading, we observed significantly reduced fluorescence signal in the root unloading zone (Figure 6F, Figure S6). This, together with our grafting experiments (Figure 6G-I), supported the role of CRK2 as a regulator of vascular loading in the source tissue. Restricted movement of not only nutrients from source to sink tissue, but also flowering signals, would explain the decreased number of leaves (Figure 4F) and altered flowering time in *crk2* (Colina et al., 2026). This could be further supported by the impact of CALSs on transport of florigens from leaves to the shoot apical meristem (Murata et al., 2025). By creating isolated developmental tissue domains (Oparka and Roberts, 2001) and by the change of those domains during sink-source transition, when an organ switches from sink to source (Roberts et al., 1997; Oparka et al., 1999), control of symplastic transport can affect the individual tissues or the destiny of whole organs. Yet, we did not observe any specific morphological alternations at organ or tissue levels in *crk2*, suggesting rather systemic effect was linked with the limitations of vasculature.

In summary, we propose CRK2 as regulator of symplastic phloem loading (Figure 7). Our study showed that CRK2 negatively regulated callose deposition *via* CALSs. Expression studies of *CALS1* and *CALS3* place the CRK2-dependent regulation of callose deposition to the vasculature, and suggested restricted vasculature loading in *crk2*. We concluded that the reduced transport of nutrients from source to sink tissues causes starch accumulation in the developmentally older leaves in *crk2*. This consequently caused starvation of sink tissues, developing leaves and roots, and led to a smaller rosette size, fewer number of leaves and short primary root length. Consistently, these phenotypes were rescued in the *crk2 cals1.5* and *crk2 cals3.1* double mutants.

**Figure 7:**
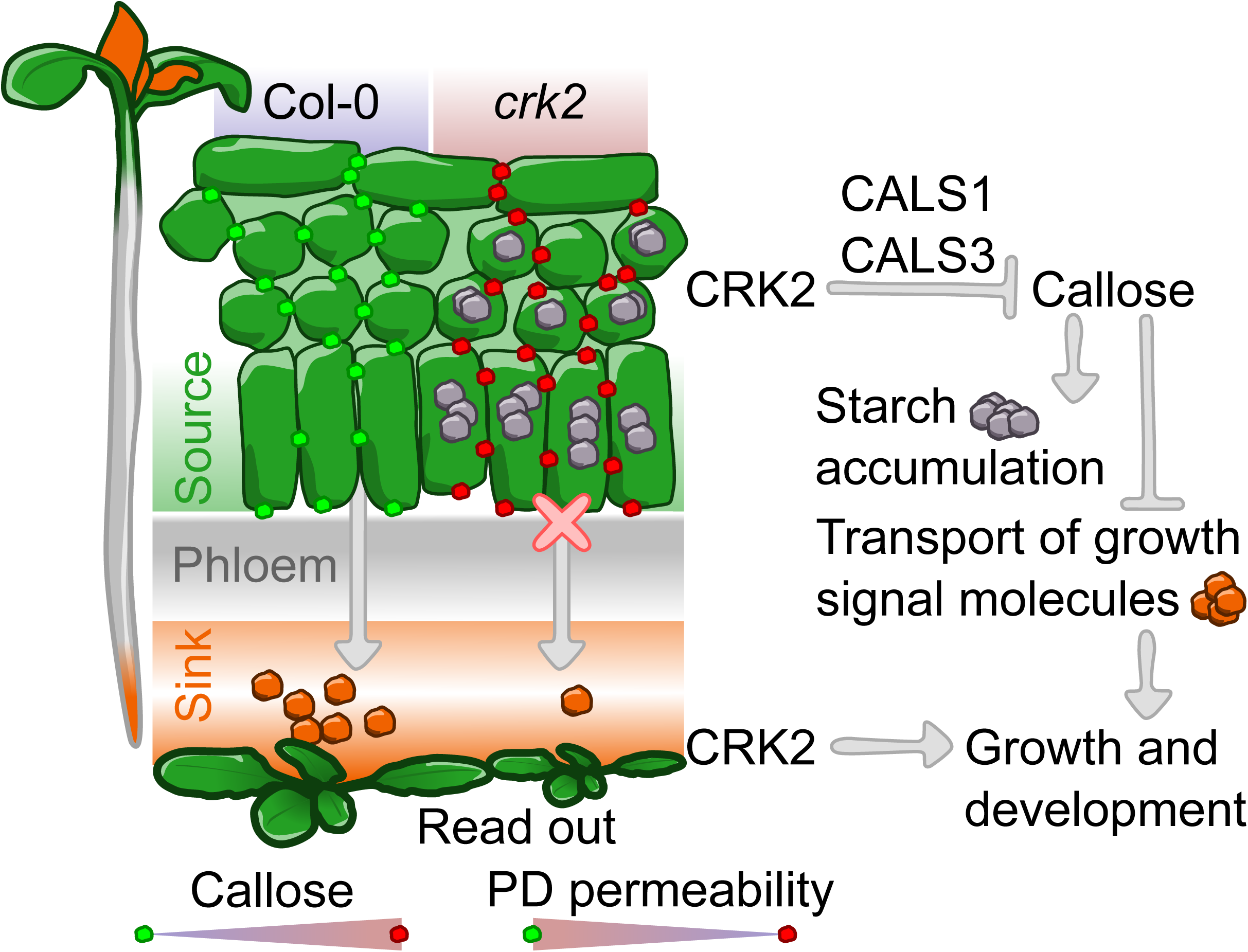
CRK2 impacts growth and developmental processes via symplastic transport regulated by CALS1 and CALS3. Mutant plants of *crk2* show increased levels of callose and starch. The presented generalized model considers the restriction of transport of growth signal molecules. This subsequently impacts growth and development. In this model, we also propose restricted transport of other macromolecules besides sugars – starch metabolism intermediates. The described accumulation could be reverted by the introduction of *cals1.5* and *cals3.1* into *crk2* mutant background. This suggests that symplastic transport, especially plasmodesmata (PD)-mediated phloem loading, regulated members of CALS proteins protein family, namely CALS1 and CALS3, play an important role in CRK2 regulated pathways.

Taken together, we propose CRK2 as a negative regulator of CALSs-mediated modulation of PD permeability, impacting growth and development of Arabidopsis plants by controlling sugar transport from source to sink tissues.

## Material and Methods

### In silico and phylogenetic analysis of proteins

The Maximum likelihood phylogenetic tree (LG model, Gamma distributed) was constructed in Mega 10 (version 10.0.5) (Kumar et al., 2018) based on a ClustalW alignment of protein sequences (Supplementary Data S1). Numbers at the nodes show bootstrap values (1000 replicates). Aligned sequences (based on alignment of twelve CALS, see above) were further processed with Mega 12 (version 12.0.9) (Kumar et al., 2018), Jalview (version 2.11.4.0) (Waterhouse et al., 2009), and molecular features of CALS1 were identified using SMART (Letunic et al., 2021) and finalized into the final figure with Inkscape (version 1.3.2).

For the projection of surface exposed phospho-sites, readily available models of CALSs were obtained from AlphaFold (Jumper et al., 2021) Protein Structure Database (https://alphafold.ebi.ac.uk/; data obtained 2025-02-27), visualized with ChimeraX (version 1.6.1; 2023-05-09) (Pettersen et al., 2021), and finalized into final figure with Inkscape (version 1.3.2).

### Molecular cloning

The encoding sequence of CALS3 N-terminal cytosolic part (CALS3_cyto_) was amplified (primers listed in Supplementary Data S3) and introduced into a bacterial expression vector (pMAL-c2X; CALS3_cyto_-pMAL-c2X). Promoter regions of *CALS1* (3000 bp) and *CALS3* (∼1500 bp), and respective 3’UTRs of *CALS1* (∼800 bp) and *CALS3* (∼670 bp) were amplified (primers listed in Supplementary Data S3) and recombined into entry (pDONR P4-P1R, pDONR P2R-P3), and, subsequently, destination (pH7m34GW) vectors (Karimi et al., 2005), using Gateway Cloning strategy (Thermo Scientific). p221a-GUS was used for the insertion of previously cloned reporter (Siligato et al., 2016). For the molecular cloning, DH5α, TOP10 or Stbl4 bacterial strains (Invitrogen) were used. Every cloned sequence was verified by the sequencing. For the agrobacterium-mediated production of transgenic lines (Clough and Bent, 1998), GV3101 pSoup (Lifeasible) strain was used. For yeast assays, the coding sequence of *CALS1* was amplified (primers listed in Supplementary Data S3) and recombined into the yeast GoldenBraid system (Pérez-González et al., 2017). Site-directed mutagenesis was used for the construction of CALS1 T19D, T21D mutations (primers listed in Supplementary Data S3). Destination vectors targeting the *Saccharomyces cerevisiae* locus *YPRC*Δ*22* and containing pTDH3-6xHA-CALS1-tTDH2 cassettes were linearized by digestion with *Not*I and transformed into the *Saccharomyces cerevisiae* BY21271 (Δ*fks1*; denotation as *fks1*) (Inoue et al., 1995) strain (NBRP Japan, *MAT*α *aro7 can1 leu2 trp1 ura3 fks1::URA3*) using the PEG3350/LiAc method (Gietz and Schiestl, 2007). Positive colonies were selected on YPD supplemented with 200 µM G418 and subsequently assayed by colony PCR (primers listed in Supplementary Data S3).

### Expression and purification of recombinant proteins

GST-CRK2_cyto_ (CRK2_cyto_-pOPINK) and GST-CRK2_cyto_KD (CRK2_cyto_KD-pOPINK, D450N) (Kimura et al., 2020) were expressed in *Escherichia coli* expression strains SHuffle T7 (New England Biolabs, C3026J). MBP (pMAL-c2X), MBP-CALS1_cyto_ (CALS1_cyto_-pOPINM) (Hunter et al., 2019), MBP-CALS3_cyto_ and MBP-FER_cyto_ (FER_cyto_-pMAL-c2X; co-transformed with pOPINK-ABI2; kindly provided by Francisco Colina and Alegría Pérez-Guillén) were expressed in BL21-CodonPlus-RIL (Agilent, 230240). Recombinant proteins were purified with glutathione sepharose 4B (GE Healthcare, 17-0756-01) or amylose resin (New England Biolabs, E8021S) beads and eluted with 10 mM maltose or 20 mM reduced glutathione, respectively.

### In vitro kinase assay

Expressed and purified recombinant proteins were incubated in ratio 1: 3-5 (kinase: substrate) 30 minutes at room temperature in reaction buffer containing γ-^32^P labelled ATP (0.2 µl/20 µl of labelled ATP with declared activity 9.25 MBq), 50 mM HEPES (pH 7.4), 1 mM DTT, 10 mM MgCl_2_, and 60 µM unlabelled ATP. Reactions were terminated by boiling the samples in SDS-based protein loading dye (98°C, 5 minutes) and proteins were separated by SDS-PAGE. Gels were stained with Coomassie brilliant blue (CBB) and dehydrated until completely dry. Dried gels were exposed to a photosensitive layer and documented with an imaging system (Molecular Dynamics, Typhoon 9410). For LC-ESI-MS/MS analysis, γ-^32^P labelled ATP was replaced by an equal amount of non-labelled ATP. After separation and staining, bands corresponding to respective recombinant proteins were excised and used for in-gel digestion.

### Protein digestion and LC-ESI-MS/MS analysis

In-gel digestion using Trypsin (Thermo Scientific, 90057), Lys-C (Promega, VA117A) and Arg-C (Promega, V188A) was performed. Digested peptide samples were dissolved in 10 µl of 0.1% (v/v) formic acid and 5 µl was injected for analysis. The LC-ESI-MS/MS analysis was performed on a nanoflow HPLC system (Thermo Scientific, Easy-nLC1000) coupled to the Q Exactive HF mass spectrometer (Thermo Scientific) equipped with a nano-electrospray ionization source. Peptides were first loaded on a trapping column and subsequently separated inline on a 15 cm C18 column (75 μm x 15 cm, ReproSil-Pur 3 µm 120 Å C18-AQ, Dr. Maisch HPLC GmbH). The mobile phase consisted of water with 0.1% (v/v) formic acid (solvent A) and acetonitrile/water (80:20 [v/v]) with 0.1% (v/v) formic acid (solvent B). Peptides were eluted using a 30 min gradient: 5% (v/v) to 21% (v/v) solvent B in 11 min, 21% (v/v) to 36% (v/v) solvent B in 9 min, and 36% (v/v) to 100% (v/v) solvent B in 5 min, followed by a 5 min wash with 100% (v/v) solvent B.

MS data was acquired automatically by using Thermo Xcalibur 4.1 software (Thermo Scientific). The data-dependent acquisition method consisted of an Orbitrap MS survey scan of mass range 350–1750 m/z with a resolution of 120000 and AGC target of 3000000. This was followed by HCD fragmentation for 10 most intense peptide ions with resolution of 15000 AGC target of 1000.

Raw files were searched for protein identifications using Proteome Discoverer 2.5 software (Thermo Scientific) connected to an in-house Mascot 2.8.2 server (Matrix Science). Data was searched against a SwissProt database (version 2022_03) with taxonomy restricted to *Arabidopsis thaliana*.

### Yeast assays

For *in vivo* CALS enzymatic activity measurements, the aniline blue-based method was employed (Janeczko, 2018; Perrine-Walker and Payne, 2021). Briefly, yeast culture was grown overnight in YPD and resuspended in milliQ water to OD_600_ = 1. Cultures were stained with 4x diluted Aniline Blue Fluorochrome (Biosupplies Australia, 100-1) in a 1:1 ratio in a microscopic chamber. The cells were imaged with confocal laser scanning microscope (Zeiss, LSM900; excitation 405 nm, detection 410-546 nm, LD LCI Plan-Apo 40x/1.2 W). Glucan levels were analysed in Fiji software with the Analyse particles tool using default settings. The mean fluorescence per cell was normalized to the *fks1* background strain in each experiment. Levels of the transgene expression were analysed using immunoblot detection after microsomal fraction isolation (see below).

### Yeast microsomal fraction isolation

Pellets from 50 ml of yeast cultures grown in YPD medium were washed with milliQ water and sedimented (5000 g, 5 min). Each culture was resuspended in homogenization buffer (50 mM HEPES, 400 mM sucrose, 100 mM KCl, 100 mM MgCl_2_, pH 7.5) and ground to fine powder using mortar and pestle in liquid nitrogen. Subsequently, the cultures were twice centrifuged (15000 g, 15 min, 4°C). The crude protein extract was then ultracentrifuged (150000 g, 60 min, 4°C) to separate the microsomal (pellet) and cytosolic (supernatant) fraction. The cytosolic fraction was decanted and the microsomal fraction resuspended in phosphate buffer (5 mM KH_2_PO_4_, 5 mM Na_2_HPO_4_, pH 7.8). For the analysis, 100 µg were used for the immunoblot. For the detection of HA-tagged protein, primary anti-HA (Invitrogen, 26183) and secondary (Promega, W4021) antibodies were used.

### Plant lines

*Arabidopsis thaliana* ecotype Col-0 was used as wild type, for the comparison with the previously described T-DNA insertion lines *crk2* (SALK_012659C) (Bourdais et al., 2015; Kimura et al., 2020), *cals1.5* (SAIL_1_H10) (Hunter et al., 2019), and *cals3.1* (SALK_068418) (Vatén et al., 2011). The presence of T-DNA insertion in a given locus was verified by PCR (used primers available in Supplementary Data S3) using Phire Plant Direct PCR Kit (Thermo Scientific, F130WH). Plant lines expressing kinase active and inactive CRK2 were described previously (Hunter et al., 2019; Kimura et al., 2020).

### Growth conditions

For *in vitro* cultivation, seeds were sterilized in 70% (v/v) ethanol with 0.02% (v/v) Triton X-100 for 5 minutes, 96% (v/v) ethanol for 1 minute and washed 5 times in deionized water and sown on half-strength Murashige and Skoog media (Duchefa, M0222.0050) pH 5.8 with 0.8% (w/v) agar, 1% (w/v) sucrose and 0.05% (w/v) MES, unless stated otherwise. Seeds were stratified in the dark, 4°C, 2 days and cultivated in a growth chamber (Sanyo, MLR-351H; 91 μmol/m^2^s^-1^) with 16 hours (23°C) light and 8 hours (18°C) dark photoperiod, unless stated otherwise. For experiments in soil, seeds were stratified in deionized water as described above, sown into soil and cultivated in growth chambers (Photon Systems Instruments, Walk-In FytoScope Walk-in growth chamber with growth units; 80 μmol/m^2^s^-1^) with 16 hours (22°C) light and 8 hours (18°C) dark – long day (LD) –, or 12 hours light (22°C) and 12 hours dark (18°C) – short day (SD) – photoperiod.

### RNA extraction, reverse transcription and qRT-PCR

7-days-old seedlings were collected and frozen in liquid nitrogen. Frozen plant material was homogenized using mortar and pestle. Homogenized material was resuspended in 800 µl of TRI reagent (Carl Roth, 9319) and centrifuged. Supernatant was mixed with 80 µl of 1-bromo-3-chloropropane (Sigma-Aldrich, B62404)and incubated at room temperature. The mixture was centrifuged and RNA precipitated with 100% (v/v) isopropyl alcohol (2:1; water phase: isopropyl alcohol). Pelleted RNA was washed with 75% (v/v) ethanol and dried. RNA was resuspended in 50µl of UltraPure DNase/RNase-Free distilled water (Invitrogen, 10977035). 1-5 µg of isolated RNA was removed from contaminating DNA with TURBO DNA-free Kit (Invitrogen, AM1907) according to the manufacturer’s protocol. Reverse transcription was done with RevertAid First Strand cDNA Synthesis Kit (Thermo Scientific, K1622), and qPCR was done using DBdirect PCR SYBR Mix SuperSens (DIANA Biotechnologies, DB-1274) following the manufacturer’s recommendation and measured with thermocycler (Bio-Rad, CFX Connect Real-Time PCR Detection System) using declared primers (primers listed in Supplementary Data S3). Relative quantification to wild type was done with the ΔΔCT method. Samples were normalized to the expression of reference gene *THIOREDOXIN-LIKE PROTEIN YLS8* (AT5G08290; Q9FE62) as described previously (Czechowski et al., 2005).

### Histochemical GUS staining

Two independent transgenic lines for each transcriptional reporter were used. 7-days-old seedlings were collected into an ice cold 90% (v/v) acetone and incubated (4°C, 60 minutes). Acetone was removed and replaced by ice cold staining buffer (5 mM NaCl, 0.01 M Tris-Cl, pH 7.2, 20% [v/v] methanol). Plans were washed 4 times with a staining buffer. Staining buffer was replaced with staining solution (5 mM NaCl, 0.01 M Tris-Cl, pH 7.2, 20% [v/v] methanol, 2 mM potassium hexacyanoferrate trihydrate, 2mM potassium ferricyanide, 1.5 mM X-GlcA cyclohexylammonium salt dissolved in dimethyl sulfoxide), and vacuum infiltrated (3 times, 5 minutes for each infiltration step). Plants were incubated (37°C, 16 hours), warmed (60°C, 15 minutes) in acidified methanol (20% [v/v] methanol, 1.5% [v/v] hydrochloric acid), and washed with 3.5 M solution of sodium hydroxide. Samples were rehydrated in decreasing ethanol gradient, saturated with 50% (v/v) glycerol, and documented with stereomicroscope (Olympus, MacroView MVX10).

### Callose analysis

Cotyledons of 7-days-old seedlings were vacuum infiltrated with 0.1% (w/v) aniline blue (Sigma-Aldrich, 415049) in phosphate-buffered saline (PBS) buffer using needleless syringe and immediately imaged with confocal laser scanning microscope (Olympus, FV3000; excitation 405 nm, detection 475-525 nm, PLAPON 60XOSC2, 8.0 μs/pixel, 12 z layers, 3.5 μm/layer). Maximum intensity projection images were subsequently processed in ImageJ (version 1.54g) and the number of PD associated deposits (stomata associated signals were masked) counted with a semi-automated approach for each manually curated field of view. For the mesophyll cells and signal associated with vasculature, cotyledons of 7-days-old seedlings were vacuum infiltrated with 0.1% (w/v) aniline blue and imaged as indicated above with the exceptions in imaging (UAPON 40XO340-2, 11 z layers, 0.740 μm/layer). Maximum intensity projection images were subsequently processed in ImageJ (version 1.54g). For mesophyll cells, the intensity of detected spots was quantified with automated approach (Brightness/contrast 20, 255; Background subtraction rolling 5; 8-bit; Threshold 20, 255; Particle analysis, size 4-300 pixel, circularity 0.20-1.00). For vasculature associated signal, signal intensity was quantified in vasculature area indicated in respective figure.

### Biolistic transformation of Arabidopsis epidermal cells

Total 5 μg of plasmid DNA in equal ratio of pB7WG2.0-SP::mRFP::KDEL and pB7WG2.0-eGFP were co-precipitated on 1.0 μm gold particles (Bio-Rad, 1652263) as described previously (Tee et al., 2022). Shoots of 7-days-old seedlings were transferred on half-strength Murashige and Skoog media (Duchefa, M0222.0050) pH 5.8 with 0.8% (w/v) agar, without sucrose and with 0.05% (w/v) MES, and positioned abaxial side up. PDS-1000/He Particle Delivery System (Bio-Rad) equipped with a 1100 psi rupture disk was used for the biolistic transformation. Transformed plants were kept in dark and analysed after 16 hours using confocal laser scanning microscope (Olympus, FV3000; GFP: excitation 488 nm, detection 500-540 nm; RFP: excitation 561 nm, detection 570-620 nm; UPLSAPO 20XO, 4.0 μs/pixel, the lowest possible number of layers to obtain image information from the whole field of view, 5 μm/layer). Numbers of cells containing GFP were counted, based on the identification of fluorescence signal at the periphery and the nucleus of the cells, and statistically analysed as previously described using median bootstrap method between *crk2* and other genotypes (Johnston and Faulkner, 2021). Statistical analysis is not indicated in the corresponding figure as we did not observe statistically significant differences.

### Phenotypic analysis

Rosettes of 21-days-old plants were excised, documented with camera (Nikon, D5600), and fresh weight was measured. Plant material was dried at 37°C until constant weight and dry weight was measured. Plates with 7-days-old seedlings were documented and primary root length between 3^rd^ and 7^th^ day measured. ImageJ (version 1.54g) was used for the respective analysis. For the vasculature analysis, 7-days-old seedlings were destained with 96% (v/v) ethanol until completely white, briefly rehydrated with deionized water and imaged with stereomicroscope (Olympus, SZX12) equipped with a camera (Promicam, 3-5CP). Obtained images were scored as indicated in the respective figure.

### Lugol’s staining

Plant material (indicated in respective section) was cultivated on half-strength Murashige and Skoog media (Duchefa, M0222.0050) pH 5.8 with 0.8% (w/v) agar, and 0.05% (w/v) MES, without sucrose in media, unless stated otherwise. 28-days-old plants were grown in a LD (see above). 7, 10, 22 and 28-days-old plants were incubated in 96% (v/v) ethanol until tissue was decolorized. Plants were rehydrated for 30 minutes in deionized water. Biological material was stained for 10 minutes in Lugol’s solution (10% [w/v] KI, 5% [w/v] I; or Sigma-Aldrich, 62650), and washed with deionized water until blue staining was distinguishable from brown background. Plants were imaged with a stereomicroscope (Olympus, MacroView MVX10), microscope (Olympus, IX83 Inverted Widefield) or camera (Nikon, D5600).

### Biochemical starch quantification

Starch was quantified from approximately 30 mg of plant material. Seedlings (indicated in respective section) were cultivated on half-strength Murashige and Skoog media (Duchefa, M0222.0050) pH 5.8 with 0.8% (w/v) agar, and 0.05% (w/v) MES, without sucrose supplementation. Seedlings were harvested at the indicated times of day into, snap frozen in liquid nitrogen and ground in a ball mill. Starch was extracted and quantified as described elsewhere (Smith and Zeeman, 2006).

### SA treatment

Seedling were sown on half-strength Murashige and Skoog media (Duchefa, M0222.0050) pH 5.8 with 0.8% (w/v) agar, and 0.05% (w/v) MES, without sucrose supplementation. 3-days-old seedlings were transplanted on growth media without sucrose supplemented with 20 µM SA (salicylic acid, 2-hydroxybenzoic acid; Sigma-Aldrich, 247588), or 4-OHBA (inactive isomer of SA, 4-Hydroxybenzoic acid; Sigma-Aldrich, H20059). 7 or 10-days-old seedlings (as indicated) were analysed. Callose deposition and starch accumulation were analysed as described previously (see above).

### Analyses of chlorophyll fluorescence kinetics (OJIP and NPQ)

4-weeks-old plants were used for the analysis with a macroscopic fluorescence imaging system with an ultrafast camera and direct imaging software for fast fluorescence transients (Küpper et al., 2019)(Photon Systems Instruments). Six plants were grown at 2 different times and were used and treated as independent repeats. Leaves were mounted in the measuring chamber and OJIP was measured by using a custom-made protocol after five minutes of leaf dark adaptation as described previously (Küpper et al., 2019), with specific conditions (100 μs shutter opening, 250 μs frame period, 4000 μmol⋅ m^-2^ s^-1^ super-saturating pulses). NPQ was measured on the same leaves after additional 5 min of dark adaptation (100 μs shutter opening, 500 μs frame period, 200 μmol⋅ m^-2^⋅ s^-1^ actinic light). Subsequently, Φ_Po_, Φ_ET2o_, and Φ_RE1o_ parameters were calculated using the Fluorcam 7 software (Photon Systems Instruments), as described previously (Stirbet and Govindjee, 2011).

### CFDA assays

6-days-old seedlings were excised in the area of hypocotyl and 1 mM CFDA (Sigma-Aldrich, 21879-25MG-F) applied in 0.8% (w/v) agar block on exposed vasculature. Plants were incubated 2 hours in darkness and imaged with confocal laser scanning microscope (Olympus, FV3000; excitation 488 nm, detection 500-600 nm, UPLSAPO 10X2, 4.0 μs/pixel, 14 z layers, 4.5 μm/layer). Z-projection images (sum slices) were subsequently processed in ImageJ (version 1.54g), by quantification of fluorescence intensity in the circular area in the zone of maximum accumulation of the fluorescent tracer.

Classical approach of CFDA assay was done as described previously (Hajný et al., 2024). Briefly, 1 mM CFDA was applied on nicked cotyledons and seedlings were imaged after 45 minutes (excitation 488 nm, emission 492-530 nm).

### Plant micro-grafting

7-day-old seedlings were used for micro-grafting and grafting assays according to a previously published method (Melnyk, 2017). For the phloem connection assay, the cotyledons were gently wounded in vascular tissues with forceps at 4 days after grafting, and CFDA was applied to the wound site. After 1 hour, phloem reconnection was considered successful when fluorescent signal was detected in the root. Fluorescence was observed using a stereo fluorescence microscope (Leica, M205 FA) equipped with a GFP filter.

### Post-grafting growth analysis

For phenotypic analysis, seedlings were transferred to half-strength Murashige and Skoog medium at 4 days after grafting and grown for an additional 10 days. The longest root was used for the measurement of the primary root length after grafting. Rosette spread area was estimated as the area enclosed by a polygon formed by connecting the outermost tips of the rosette leaves.

### Statistical analysis

Number of experimental repeats and the balanced number of individual plants are stated in respective sections. For the statistical analyses, the integrated development environment RStudio (2023-12-1; Build 402; Posit Software) for R language (2024-04-24, 4.4.0; R Foundation) was used, together with ggplot2 (3.5.1) (Wickham, 2016), agricolae (1.3-7) (Felipe de Mendiburu, 2006), and dplyr (1.1.4) (Wickham et al., 2023) packages. The specific statistical tests used for experiments are listed in the corresponding figure legends.

### Gene IDs

CALS1 (AT1G05570; Q9AUE0), CALS3 (AT5G13000; Q9LXT9), CRK2 (AT1G70520; Q9CAL3)

## Acknowledgements

The project was supported by the Czech Science Foundation Grantová Agentura České Republiky grants 23-04866S and 25-15633S (to MW)) and 22-35680M (to RP), the Palacký University Young Researcher Grant JG_2024_003 implemented within (to JH), GA UK nr. 270823 (DU), and by a grant from the Ministry of Education, Youth and Sports (MEYS) of the Czech Republic with co-financing from the EU (CZ.02.1.01/0.0/0.0/15_003/0000336, “KOROLID” to HK). YS and CWM were supported by a Horizon Europe ERC Consolidator Grant (101126239 GRAFT-ABLE). Part of the work was carried out with the support of a Growth Facility (Biology Centre CAS Core Facilities; IPMB Biology Centre CAS). We thank the Laboratory of Microscopy and Histology of the Institute of Entomology, Mr. Jan Kadlec (Growth Facility, Biology Centre CAS Core Facilities), the Media Kitchen of Institute of Plant Molecular Biology at the Biology Centre CAS for support and provision of instruments, and Dr. Iva Mozgová (Biology Centre CAS, Czech Republic) for the sharing of phenotyping infrastructure. Mass spectrometry analyses were performed at the Turku Proteomics Facility supported by Biocenter Finland. Microscopy was performed at the Imaging Facility of the IEB CAS, supported by MEYS CR LM2023050 ‘Czech-BioImaging’ and IEB CAS (Prague). We would like to also thank to Dr. Christine Faulkner (John Innes Centre, United Kingdom) for the insightful advices and vectors for bombardment experiments, Dr. Lin Xi (University of Hohenheim, Germany) for suggestions about CFDA assay improvements, Ms. Nerea Valdebenito Alamar (University of Helsinki, Finland) for participating in the molecular cloning of the N-terminal part of the CALS3 coding sequence, and Dr. María Illescas Morente for critical reading of manuscript, and together with Dr. Francisco Colina and Ms. Alegría Pérez-Guillén (Biology Centre CAS, Czech Republic) for the kind provision of FER_cyto_-pMAL-c2X and respective recombinant protein.

## Author contributions

AZ, JK-W and MW conceived, conceptualized and designed the experiments. MW, RP, DU, JH, HK, and CWM acquired research grants. AZ performed *in silico* analyses, *in vitro* kinase assay, expression analyses, bombardment, starch staining and statistical analyses. AZ and MS performed phenotypic analyses; MP and JM carried out phosphoproteomics analyses; DU and RP carried out yeast assays, JK-W carried out biochemical quantification of starch; AZ and SJ carried out callose staining; FM, EA, and HK analysed chlorophyll fluorescence kinetics; AZ and JH performed CFDA assays. YS and CWM did grafting experiments. AZ wrote the initial draft; JK-W, SJ and MW edited and reviewed the manuscript. All the authors commented on the manuscript and approved the final version.

## Data availability

Data used in this study are available as supplementary data. Materials used for the research are available from the corresponding author upon request.

---

**Supplementary Figure S1: Analysis of CALS1 and CALS3 phosphorylation.** (A) Aligned N-terminal regions of CALS1 and CALS3, which were also used for the construction of the phylogeny tree, showed striking conservation of potential phospho-sites (red), namely those in annotated domains Vta1 (VPS20-associated protein 1) and FKS1 (1,3-beta-glucan synthase domain). Phospho-sites indicated in PhosPhAt 4.0 database (green circle). CRK2-specific phosphorylation sites identified using *in vitro* phosphoproteomics (grey asterisk). The presence of additional symbols indicates identification in the independent experiments. LCR – low complexity region, TMD1 – first N-terminal transmembrane domain of CALS. (B) Predicted structures of CALS1 and CALS3 support the proposed phosphorylation-based regulation as significant number of potential phospho-sites (red) on N-terminal region (grey) is surface-exposed. Identified CRK2-dependent phospho-sites are indicated. (C) Cytoplasmic region of CRK2 containing active kinase domain (CRK2_cyto_) shows higher activity compared to cytoplasmic region of FERONIA (FER) containing active kinase domain (FER_cyto_) (left panel). *In vitro* phosphorylation of N-terminal regions of CALS1 (CALS1_cyto_) and CALS3 (CALS3_cyto_) was not visible even after differential exposure of CRK2 (middle panel) and FER (right panel) assays. CBB – Coomassie brilliant blue, MBP – maltose binding protein, GST – glutathione S-transferase, relative molecular weight is in kDa. (D) Alignment of CALS sites identified as phosphorylated by CRK2 (highlighted). Number indicates position in the sequence of respective CALS. (E) Expression of CALS_WT_ and CALS1_mut_ in tested yeasts is comparable. CF – cytosolic fraction, MF – microsomal fraction, relative molecular weight is in kDa.

**Supplementary Figure S2:** (A) PCR-based selection of double mutants *crk2 cals3.1* and *crk2 cals3.1* lines from previously described *crk2*, *cals1.5* and *cals3.1* prenatal T-DNA insertional knock-out lines. Blank – extraction buffer replacing DNA sample. (B) Representative image of the analyses of callose in mesophyll cells (Meso) and vasculature (Vasc), as indicated by the area marked by white dashed line. Bar – 100 µm. (C) Quantification of aniline blue fluorescence intensity in mesophyll cells. Different presentation of data was selected because of number of datapoints used in the analysis. Kruskal–Wallis test (Df = 5, H = 25.931) with Wilcoxon test. Cross indicates mean, blue dashed line indicates mean value of Col-0, n ≥ 2900. (D) Quantification of fluorescence signal intensity associated with the vasculature, as indicated in B. Kruskal–Wallis test (Df = 5, H = 12.553) with Wilcoxon test. Different colours indicate independent experiments, horizontal line indicates median, cross indicates mean, whiskers show minimum and maximum value, black dots indicate outliers, box indicates upper and lower quartiles, n = 36. (C, D) Different letters indicate statistical significance p < 0.05.

**Supplementary Figure S3: Selected double mutant plants showed similar reversion of rosette area (A, C) and dry weight (B, D). This reversion is not altered by the different photoperiod (A, B – long day, LD; C, D – short day, SD).** (A) Kruskal–Wallis test (Df = 5, H = 61.801) with Wilcoxon test. (B) One-way ANOVA (Df = 5, 186; F = 7.272) with Tukey’s HSD. (C) Kruskal–Wallis test (Df = 5, H = 76.12) with Wilcoxon test. (D) Kruskal–Wallis test (Df = 5, H = 62.209) with Wilcoxon test. (E, F) Normalised rosette area (E) and dry weight (F).(E) Wilcoxon test (W = 927). (F) Wilcoxon test (W = 863). (G) Representative pictures of 7-days-old seedling grown (LD) on indicated concentration (% [w/v]) supplemented growth media. (H, I, J) Supplementation with sucrose (% [w/v]) in media reverted small rosette area (I) and reduced true leaf number (J) phenotypes. CRK2 – *pCRK2::CRK2-mVenus* #1-22/*crk2*, CRK2^D450N^ – *pCRK2::CRK2^D450N^-mVenus*/*crk2.* (I) Two-Way ANOVA (treatment: Df = 2, 268, F = 15.363; genotype: Df = 3, 268, F = 123.751; interaction: Df = 6,268, F = 2.551). (J) Two-Way ANOVA (treatment: Df = 2, 268, F = 8.715; genotype: Df = 3, 268, F = 163.526; interaction: Df = 6, 268, F = 2.770). (I, J) with Tukey’s HSD. (E, F, I, J) Values normalized to the mean value of Col-0 of the respective repeat and treatment. (G, H) Bar – 10 mm. (A, B, C, D, E, F, I, J) Different colours indicate independent experiments, horizontal line indicates median, cross indicates mean, whiskers show minimum and maximum value, black dots indicate outliers, box indicates upper and lower quartiles. Different letters indicate statistical significance p < 0.05, n < 17.

**Supplementary Figure S4: *crk2* accumulated starch in various developmental stages.** (A) Lugol’s staining of 7-days-old and (B) 10-days-old seedlings at the end of the day (EoD) and end of the night (EoN). (A, B) Seedlings grew on media containing 1% (w/v) sucrose. (C) Lugol’s staining of 28-days-old plants. (D) Biochemical quantification of starch in 21-days-old plants grown on media supplemented with sucrose (suc). C – cotyledon, number indicates respective leaf. Two-Way ANOVA (treatment: Df = 2, 147, F = 24.488; genotype: Df = 6, 147, F = 848.142; interaction: Df = 12, 147, F = 6.504) with Tukey’s HSD, n = 8. (E) Lugo’s staining of the root tips of *crk2* and complementation line with kinase inactive CRK2 (CRK2^D450N^, *pCRK2::CRK2^D450N^-mVenus*/*crk2*) and kinase active CRK2 (CRK2, *pCRK2::CRK2-mVenus* #1-22/*crk2*). Bar – 20 μm. (F) Lugol’s staining and (G) biochemical quantification of starch after application of salicylic acid (SA), where inactive isomer of SA – 4-hydroxybenzoic acid (4-OHBA) – was used as negative control. (G) Two-Way ANOVA (treatment: Df = 1, 44, F = 15.172; genotype: Df = 1, 44, F = 87.022; interaction: Df = 1, 44, F = 5.715) with Tukey’s HSD, n = 12. (H) Elevation of starch was also accompanied by statistically insignificantly elevated levels of callose. Two-Way ANOVA (treatment: Df = 1, 92, F = 1.590; genotype: Df = 1, 92, F = 12.068; interaction: Df = 1, 92, F = 0.017) with Tukey’s HSD, n = 24. (D, G, H) Different colours indicate independent experiments, horizontal line indicates median, cross indicates mean, whiskers show minimum and maximum value, black dots indicate outliers, box indicates upper and lower quartiles. Different letters indicate statistical significance p < 0.05. (A, B, C, E, F) Bar – 10 mm.

**Supplementary Figure S5: Analysis of non-photochemical quenching (NPQ).** i – irradiance, r – relaxation; Kruskal–Wallis test followed by Wilcoxon test; i1 (Df = 3, H = 4.2214), i2 (Df = 3; H = 5.0888), i3 (Df = 3, H = 2.203), i4 (Df = 3, H = 2.194), r1 (Df = 3, H = 11.467), r2 (Df = 3; H = 12.304), r3 (Df = 3; H = 16.576). Different colours indicate independent experiments, horizontal line indicates median, cross indicates mean, whiskers show minimum and maximum value, black dots indicate outliers, box indicates upper and lower quartiles. Different letters indicate statistical significance p < 0.05, n = 12.

**Supplementary Figure S6: Vasculature of *crk2* is comparable with wild type.** (A) Vasculature patterning in cotyledons. Cotyledon patterning was scored into the following classes as indicated: Ex – expected pattern (pale grey), Pa – at least one partially developed secondary vein (dark grey), Ab – absence of at least one secondary vein (white), Co – combination of Pa and Ab (black). χ^2^ test (Df = 3, χ^2^ = 2.558). Different letters indicate statistical significance, n ≥ 58. (B) We did not observe any pronounced differences in root architecture. (C) Analysis of root meristem length. Welch’s two sample t-test (Df = 35.695, t = 1.0558). (D, E) Classical CFDA assay testing impact of the genetic interaction. We observed less fluorescence signal in *crk2* while independent introduction of *CALS1* or *CALS3* mutant alleles caused increased observed fluorescent signal in the root tip. (B, D) Bar – 100 µm. (E) Quantification of fluorescence intensity in the root tip. Kruskal–Wallis test (Df = 4, H = 24.472) with Wilcoxon test. (C, E) Different letters indicate statistical significance p < 0.05, n ≥ 20. Horizontal line indicates median, cross indicates mean, whiskers show minimum and maximum value, black dots indicate outliers, box indicates upper and lower quartiles. (D, E) Comparison of wild type and *crk2* is in Figure 6, wild type plants were not analysed to the reduce number of analysed samples. (F) Phloem reconnection rate of wild type plants and *crk2*. Percentage of plants (labelled as shoot/root) showing successful reconnection (number of positive plants/number of tested plants). (C, D, F) Different colours indicate independent experiments.

**Supplementary Data S1: Multiple sequence alignment used for the construction of phylogenetic tree.**

**Supplementary Data S2: List of identified CRK2-dependent phosphorylation sites in this study.**

**Supplementary Data S3: List of used primers in this study.**

## Notes

### Competing Interest Statement

The authors have declared no competing interest.

### Summary of Updates

We have added experimental evidence for the regulation of callose synthase activity by phosphorylation, grafting experiments as well as additional sugar measurements and callose quantifications.

