## Supplementary Figures for "CYSTEINE-RICH RLK2 regulates development *via* callose synthase-dependent symplastic transport in *Arabidopsis*"

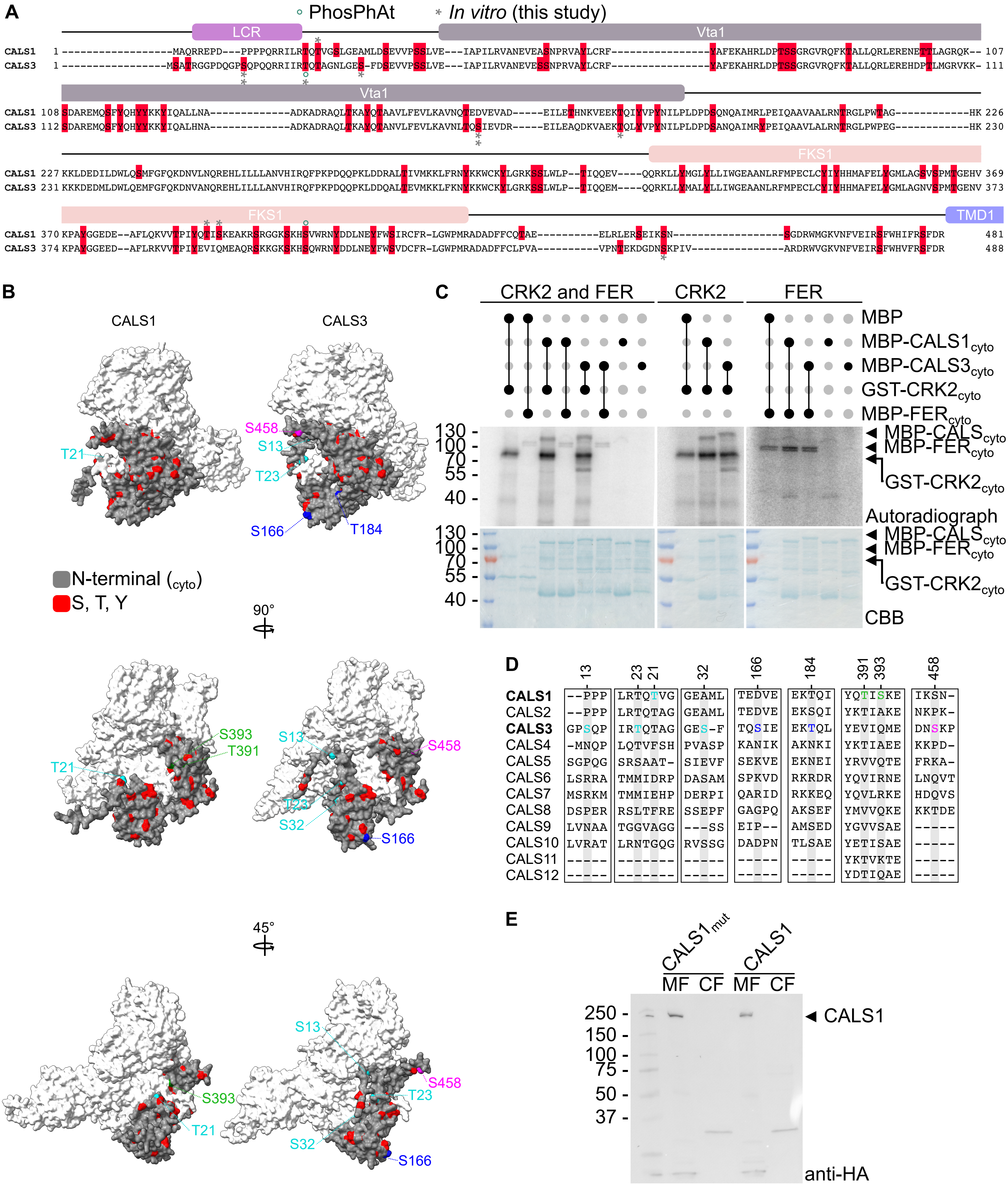
A
PhosPhAt
In vitro (this study)
LCR
Vta1
FKS1
TMD1
CALS1
CALS3
B
CALS1
CALS3
T21
S458
S13
T23
S166
T184
N-terminal (cyto)
S, T, Y
90°
S393
T391
S32
45°
C
CRK2 and FER
CRK2
FER
MBP
MBP-CALS1cyto
MBP-CALS3cyto
GST-CRK2cyto
MBP-FERcyto
130
100
70
55
40
MBP-CALScyto
Autoradiograph
CBB
D
13
23
21
32
166
184
391
393
458
CALS1 --PPP LRTQTVG GEAML TEDVE EKTQI YQTISKE IKSN-
CALS2 --PPP LRTQTAG GEAML TEDVE EKSQI YKTIAKE NKPK-
CALS3 GPSQP IRTQTAG GES-F TQSIE EKTQL YEVIQME DNSKP
CALS4 -MNQP LQTVFSH PVASP KANIK AKNKI YKTIAEE KKPD-
CALS5 SGPQG SRSAAT- SIEVF SEKVE EKNEI YRVVQTE FRKA-
CALS6 LSRRA TMMIDRP DASAM SPKVD RKRDR YQVIRNE LNQVT
CALS7 MSRKM TMMIEHP DERPI QARID RKKEQ YQVLRKE HDQVS
CALS8 DSPER RSLTFRE SSEPF GAGPQ AKSEF YMVVQKE KKTDE
CALS9 LVNAA TGGVAGG ---SS EIP-- AMSED YGVVSAE -----
CALS10 LVRAT LRNTGQG RVSSG DADPN TLSAE YETISAE -----
CALS11 ----- ------- ----- ----- ----- YKTVKTE -----
CALS12 ----- ------- ----- ----- ----- YDTIQAE -----
E
CALS1mut
CALS1
MF
CF
250
150
100
75
50
37
CALS1
anti-HA

**Supplementary Figure S1: Analysis of CALS1 and CALS3 phosphorylation.** (A) Aligned N-terminal regions of CALS1 and CALS3, which were also used for the construction of the phylogeny tree, showed striking conservation of potential phospho-sites (red), namely those in annotated domains Vta1 (VPS20-associated protein 1) and FKS1 (1,3-beta-glucan synthase domain). Phospho-sites indicated in PhosPhAt 4.0 database (green circle). CRK2-specific phosphorylation sites identified using *in vitro* phosphoproteomics (grey asterisk). The presence of additional symbols indicates identification in the independent experiments. LCR – low complexity region, TMD1 – first N-terminal transmembrane domain of CALS. (B) Predicted structures of CALS1 and CALS3 support the proposed phosphorylation-based regulation as significant number of potential phospho-sites (red) on N-terminal region (grey) is surface-exposed. Identified CRK2-dependent phospho-sites are indicated. (C) Cytoplasmic region of CRK2 containing active kinase domain ($CRK2_{cyto}$) shows higher activity compared to cytoplasmic region of FERONIA (FER) containing active kinase domain ($FER_{cyto}$) (left panel). *In vitro* phosphorylation of N-terminal regions of CALS1 ($CALS1_{cyto}$) and CALS3 ($CALS3_{cyto}$) was not visible even after differential exposure of CRK2 (middle panel) and FER (right panel) assays. CBB – Coomassie brilliant blue, MBP – maltose binding protein, GST – glutathione S-transferase, relative molecular weight is in kDa. (D) Alignment of CALS sites identified as phosphorylated by CRK2 (highlighted). Number indicates position in the sequence of respective CALS. (E) Expression of $CALS_{WT}$ and $CALS1_{mut}$ in tested yeasts is comparable. CF – cytosolic fraction, MF – microsomal fraction, relative molecular weight is in kDa.

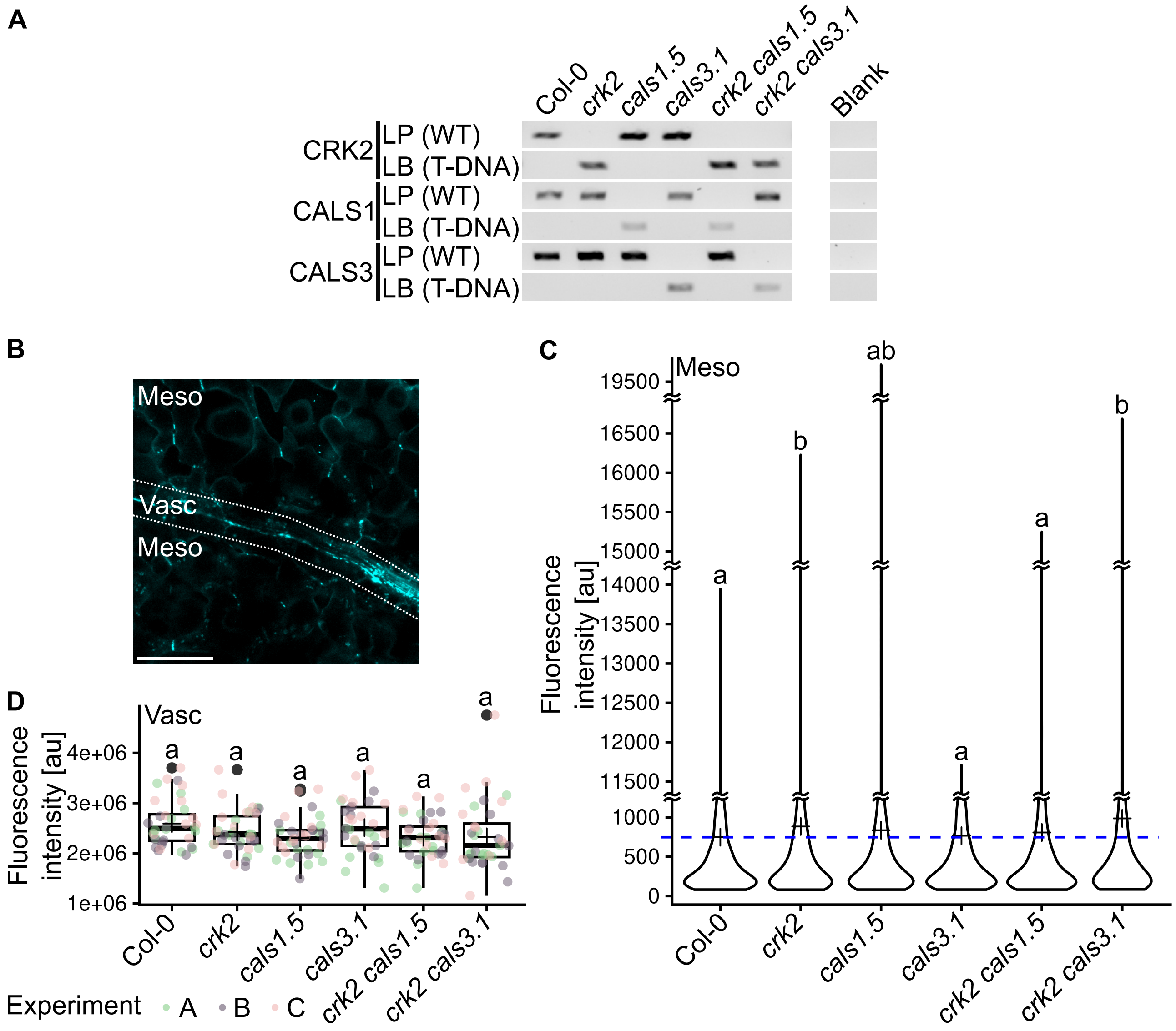


**Supplementary Figure S2:** (A) PCR-based selection of double mutants *crk2 cals3.1* and *crk2 cals3.1* lines from previously described *crk2*, *cals1.5* and *cals3.1* prenatal T-DNA insertional knock-out lines. Blank – extraction buffer replacing DNA sample. (B) Representative image of the analyses of callose in mesophyll cells (Meso) and vasculature (Vasc), as indicated by the area marked by white dashed line. Bar – 100 µm. (C) Quantification of aniline blue fluorescence intensity in mesophyll cells. Different presentation of data was selected because of number of datapoints used in the analysis. Kruskal–Wallis test (Df = 5, H = 25.931) with Wilcoxon test. Cross indicates mean, blue dashed line indicates mean value of Col-0, $n \geq 2900$. (D) Quantification of fluorescence signal intensity associated with the vasculature, as indicated in B. Kruskal–Wallis test (Df = 5, H = 12.553) with Wilcoxon test. Different colours indicate independent experiments, horizontal line indicates median, cross indicates mean, whiskers show minimum and maximum value, black dots indicate outliers, box indicates upper and lower quartiles, n = 36. (C, D) Different letters indicate statistical significance $p < 0.05$.

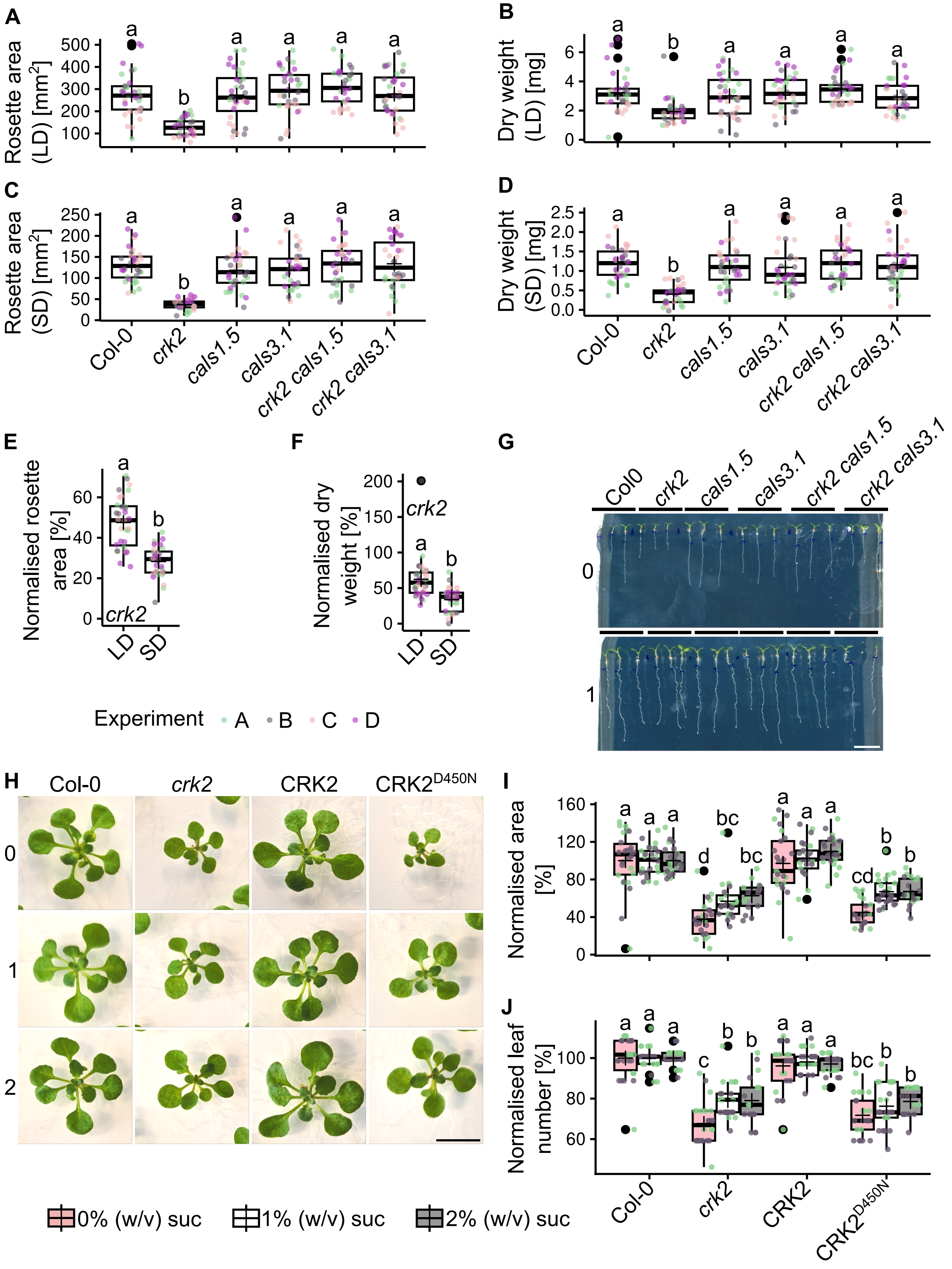
A
Rosette area (LD) [mm²]
B
Dry weight (LD) [mg]
C
Rosette area (SD) [mm²]
D
Dry weight (SD) [mg]
Col-0
crk2
cals1.5
cals3.1
crk2 cals1.5
crk2 cals3.1
E
Normalised rosette area [%]
F
Normalised dry weight [%]
LD
SD
G
Col0
Experiment A B C D
H
CRK2
CRK2D450N
I
Normalised area [%]
J
Normalised leaf number [%]
0% (w/v) suc
1% (w/v) suc
2% (w/v) suc

**Supplementary Figure S3: Selected double mutant plants showed similar reversion of rosette area (A, C) and dry weight (B, D). This reversion is not altered by the different photoperiod (A, B – long day, LD; C, D – short day, SD).** (A) Kruskal–Wallis test (Df = 5, H = 61.801) with Wilcoxon test. (B) One-way ANOVA (Df = 5, 186; F = 7.272) with Tukey's HSD. (C) Kruskal–Wallis test (Df = 5, H = 76.12) with Wilcoxon test. (D) Kruskal–Wallis test (Df = 5, H = 62.209) with Wilcoxon test. (E, F) Normalised rosette area (E) and dry weight (F).(E) Wilcoxon test (W = 927). (F) Wilcoxon test (W = 863). (G) Representative pictures of 7-days-old seedling grown (LD) on indicated concentration (% [w/v]) supplemented growth media. (H, I, J) Supplementation with sucrose (% [w/v]) in media reverted small rosette area (I) and reduced true leaf number (J) phenotypes. CRK2 – *pCRK2::CRK2-mVenus #1-22/crk2*, CRK2D450N – *pCRK2::CRK2D450N-mVenus/crk2*. (I) Two-Way ANOVA (treatment: Df = 2, 268, F = 15.363; genotype: Df = 3, 268, F = 123.751; interaction: Df = 6, 268, F = 2.551). (J) Two-Way ANOVA (treatment: Df = 2, 268, F = 8.715; genotype: Df = 3, 268, F = 163.526; interaction: Df = 6, 268, F = 2.770). (I, J) with Tukey's HSD. (E, F, I, J) Values normalized to the mean value of Col-0 of the respective repeat and treatment. (G, H) Bar – 10 mm. (A, B, C, D, E, F, I, J) Different colours indicate independent experiments, horizontal line indicates median, cross indicates mean, whiskers show minimum and maximum value, black dots indicate outliers, box indicates upper and lower quartiles. Different letters indicate statistical significance $p < 0.05$, $n < 17$.

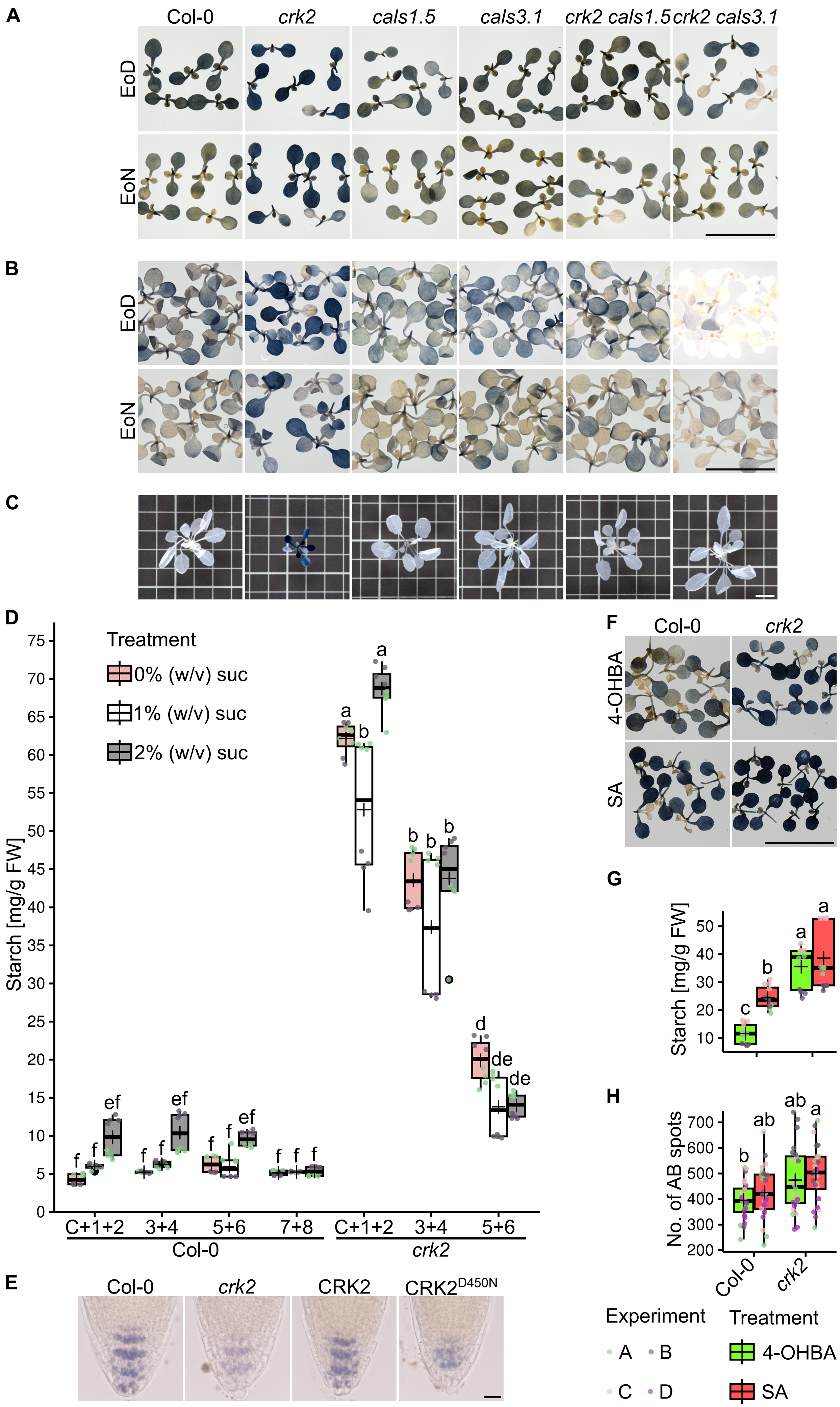
A
Col-0
crk2
cals1.5
cals3.1
crk2 cals1.5
crk2 cals3.1
EoD
EoN
B
EoD
EoN
C
D
Treatment
0% (w/v) suc
1% (w/v) suc
2% (w/v) suc
Starch [mg/g FW]
C+1+2
3+4
5+6
7+8
C+1+2
3+4
5+6
Col-0
crk2
E
Col-0
crk2
CRK2
CRK2D450N
F
Col-0
crk2
4-OHBA
SA
G
Starch [mg/g FW]
H
No. of AB spots
Col-0
crk2
Experiment
A
B
C
D
Treatment
4-OHBA
SA

**Supplementary Figure S4: *crk2* accumulated starch in various developmental stages.** (A) Lugol's staining of 7-days-old and (B) 10-days-old seedlings at the end of the day (EoD) and end of the night (EoN). (A, B) Seedlings grew on media containing 1% (w/v) sucrose. (C) Lugol's staining of 28-days-old plants. (D) Biochemical quantification of starch in 21-days-old plants grown on media supplemented with sucrose (suc). C – cotyledon, number indicates respective leaf. Two-Way ANOVA (treatment: Df = 2, 147, F = 24.488; genotype: Df = 6, 147, F = 848.142; interaction: Df = 12, 147, F = 6.504) with Tukey's HSD, n = 8. (E) Lugo's staining of the root tips of crk2 and complementation line with kinase inactive CRK2 (CRK2D450N, *pCRK2::CRK2D450N-mVenus/crk2*) and kinase active CRK2 (CRK2, *pCRK2::CRK2-mVenus #1-22/crk2*). Bar – 20 µm. (F) Lugol's staining and (G) biochemical quantification of starch after application of salicylic acid (SA), where inactive isomer of SA – 4-hydroxybenzoic acid (4-OHBA) – was used as negative control. (G) Two-Way ANOVA (treatment: Df = 1, 44, F = 15.172; genotype: Df = 1, 44, F = 87.022; interaction: Df = 1, 44, F = 5.715) with Tukey's HSD, n = 12. (H) Elevation of starch was also accompanied by statistically insignificantly elevated levels of callose. Two-Way ANOVA (treatment: Df = 1, 92, F = 1.590; genotype: Df = 1, 92, F = 12.068; interaction: Df = 1, 92, F = 0.017) with Tukey's HSD, n = 24. (D, G, H) Different colours indicate independent experiments, horizontal line indicates median, cross indicates mean, whiskers show minimum and maximum value, black dots indicate outliers, box indicates upper and lower quartiles. Different letters indicate statistical significance $p < 0.05$. (A, B, C, E, F) Bar – 10 mm.

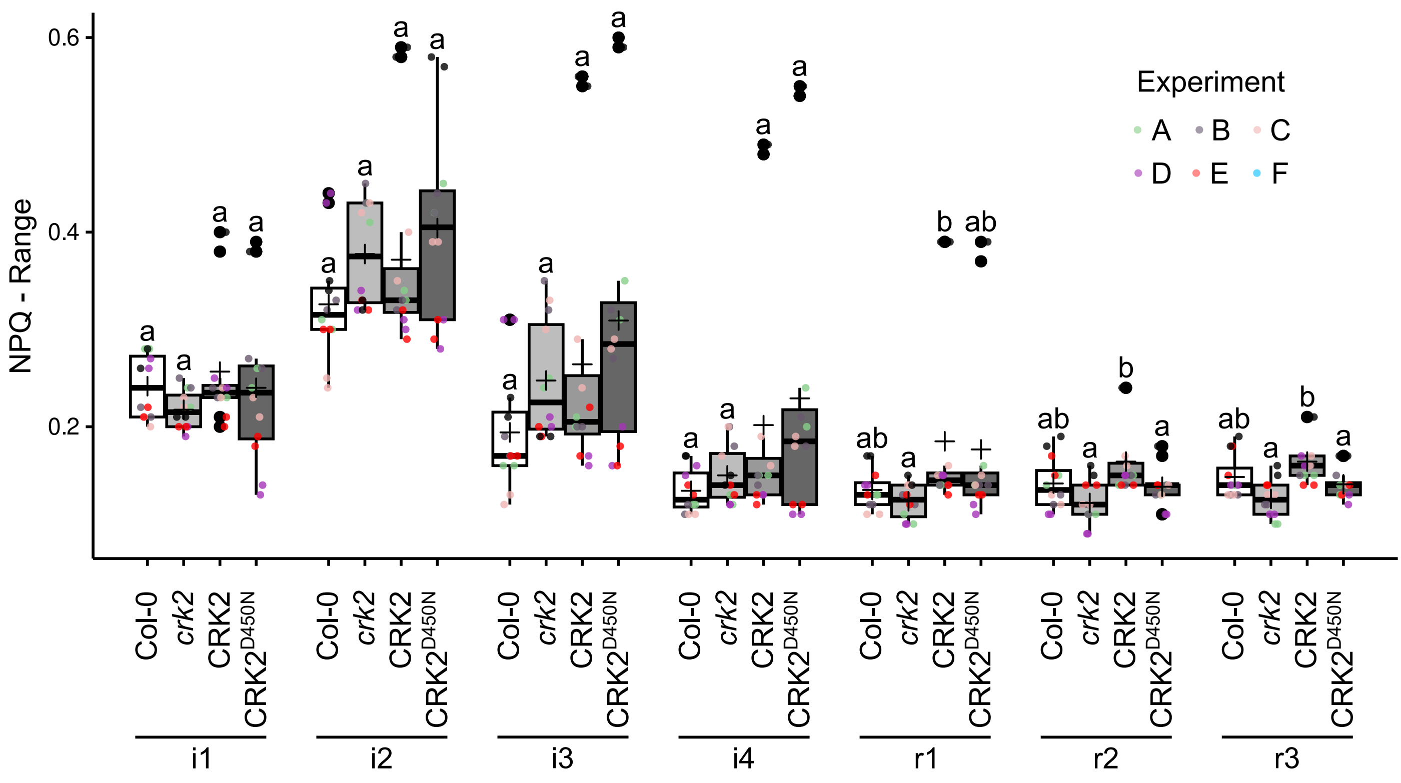


**Supplementary Figure S5: Analysis of non-photochemical quenching (NPQ)**. i – irradiance, r – relaxation; Kruskal–Wallis test followed by Wilcoxon test; i1 (Df = 3, H = 4.2214), i2 (Df = 3; H = 5.0888), i3 (Df = 3, H = 2.203), i4 (Df = 3, H = 2.194), r1 (Df = 3, H = 11.467), r2 (Df = 3; H = 12.304), r3 (Df = 3; H = 16.576). Different colours indicate independent experiments, horizontal line indicates median, cross indicates mean, whiskers show minimum and maximum value, black dots indicate outliers, box indicates upper and lower quartiles. Different letters indicate statistical significance $p < 0.05$, n = 12.

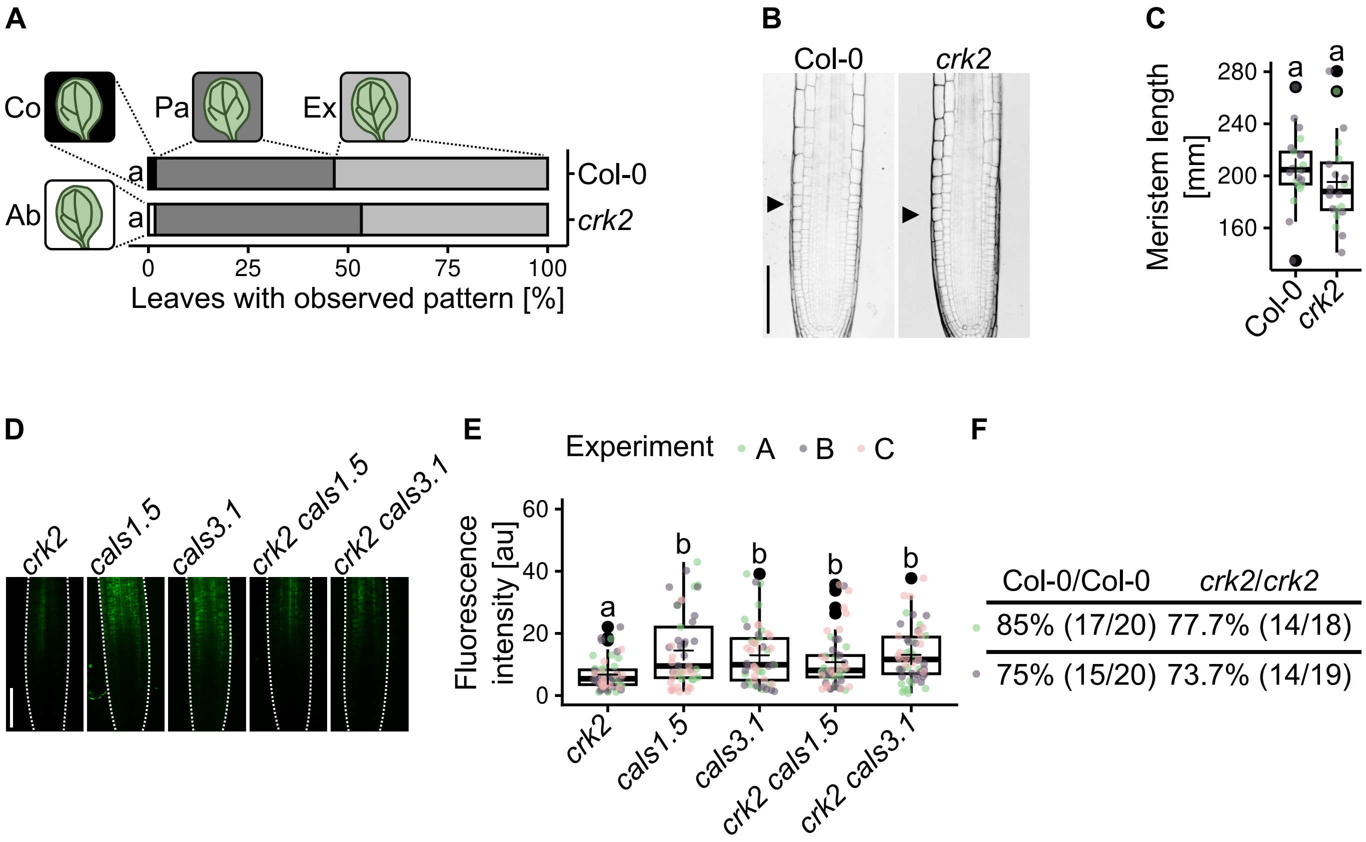


**Supplementary Figure S6: Vasculature of *crk2* is comparable with wild type.** (A) Vasculature patterning in cotyledons. Cotyledon patterning was scored into the following classes as indicated: Ex – expected pattern (pale grey), Pa – at least one partially developed secondary vein (dark grey), Ab – absence of at least one secondary vein (white), Co – combination of Pa and Ab (black). $\chi^2$ test (Df = 3, $\chi^2$ = 2.558). Different letters indicate statistical significance, n ≥ 58. (B) We did not observe any pronounced differences in root architecture. (C) Analysis of root meristem length. Welch's two sample t-test (Df = 35.695, t = 1.0558). (D, E) Classical CFDA assay testing impact of the genetic interaction. We observed less fluorescence signal in *crk2* while independent introduction of *CALS1* or *CALS3* mutant alleles caused increased observed fluorescent signal in the root tip. (B, D) Bar – 100 µm. (E) Quantification of fluorescence intensity in the root tip. Kruskal–Wallis test (Df = 4, H = 24.472) with Wilcoxon test. (C, E) Different letters indicate statistical significance $p < 0.05$, n ≥ 20. Horizontal line indicates median, cross indicates mean, whiskers show minimum and maximum value, black dots indicate outliers, box indicates upper and lower quartiles. (D, E) Comparison of wild type and *crk2* is in Figure 6, wild type plants were not analysed to the reduce number of analysed samples. (F) Phloem reconnection rate of wild type plants and *crk2*. Percentage of plants (labelled as shoot/root) showing successful reconnection (number of positive plants/number of tested plants). (C, D, F) Different colours indicate independent experiments.
